# Elucidating lipid nanoparticle properties and structure through biophysical analyses

**DOI:** 10.1101/2024.12.19.629496

**Authors:** Marshall S. Padilla, Sarah J. Shepherd, Andrew R. Hanna, Martin Kurnik, Xujun Zhang, Michelle Chen, James Byrnes, Ryann A. Joseph, Hannah M. Yamagata, Adele S. Ricciardi, Kaitlin Mrksich, David Issadore, Kushol Gupta, Michael J. Mitchell

## Abstract

Designing lipid nanoparticle (LNP) delivery systems with specific targeting, potency and minimal side effects is crucial for their clinical use. However, traditional characterization methods, such as dynamic light scattering, cannot accurately quantify physicochemical properties of LNPs and how they are influenced by the lipid composition and mixing method. Here we structurally characterize polydisperse LNP formulations by applying emerging solution-based biophysical methods that have higher resolution and provide biophysical data beyond size and polydispersity. These techniques include sedimentation velocity analytical ultracentrifugation, field-flow fractionation followed by multi-angle light scattering, and size-exclusion chromatography in-line with synchrotron small-angle X-ray scattering. We show that LNPs have intrinsic polydispersity in size, RNA loading, and shape, which depends on both the formulation technique and lipid composition. Lastly, we predict LNP transfection in vitro and in vivo by examining the relationship between mRNA translation and physicochemical characteristics. Solution-based biophysical methods will be essential for determining LNP structure-function relationships, facilitating the creation of new design rules for LNPs.

## Introduction

Lipid nanoparticles (LNPs) have emerged as the preeminent non-viral drug delivery vehicle for nucleic acids, such as messenger RNA (mRNA), and are utilized for vaccines, protein replacement therapies, and gene therapy^1–3^. Their success has been demonstrated by the U.S. Food and Drug Administration’s (FDA) approval of an siRNA LNP therapeutic, Alnylam Pharmaceutical’s Onpattro^TM^, and the FDA approval of two COVID-19 mRNA LNP vaccines, Moderna’s Spikevax^TM^ and Pfizer/BioNTech’s Comirnaty^TM4^. LNPs typically consist of four lipid components: an ionizable lipid, a phospholipid, a cholesterol, and a lipid-anchored PEG, which all contribute to LNP structure, size, and RNA encapsulation^5^. The formulation of LNPs occurs via rapid mixing of an ethanol phase containing the lipids and a low pH aqueous phase containing RNA. Changes in lipid composition and production methods can substantially impact LNP structure and physiochemical properties, which can ultimately determine LNP stability, performance, and biodistribution^6^. Previous studies have demonstrated that microfluidic mixing devices, such as those incorporating staggered herringbone micromixer architectures, can improve control over desirable LNP physical properties^7,8^. Furthermore, the laminar flow conditions employed in these microfluidic devices enables a rapid, controlled, and reproducible fluidic profile where microfluidic-generated LNPs can be significantly more potent than bulk mixed LNPs for mRNA delivery^9^.

Although novel formulation modalities can enhance the efficacy of LNPs, understanding the mechanism underlying these improvements is hindered by the current characterization practices, which suffer from low sensitivity and resolution^10^. For example, many traditional techniques, such as the Quant-it^TM^ RiboGreen RNA assay, cannot differentiate between LNPs containing no RNA and RNA-containing LNPs; thus, it is difficult to accurately describe RNA loading heterogeneity^11^. This is essential as recent studies have found that up to 80% of LNPs are empty^12–14^. Dynamic light scattering (DLS), a common technique to characterize LNP size distributions, also cannot differentiate subpopulations, cannot discern asphericity, has limited resolution for polydisperse samples, and is biased towards larger species. Similarly, cryo-TEM is unable to distinguish loaded and unloaded LNPs and may not accurately describe LNP size, morphology, and structure since LNPs are evaluated in a vitrified state prepared onto a grid that is not representative of LNPs in solution.

Hence, current efforts to define relationships between the biological efficacy of a formulation with the individual lipid structures, excipient ratios, formulation technique, and physiochemical characteristics have yielded weak and conflicting correlations. While several studies have emerged in the past five years employing more advanced characterization strategies, they are focused on a single technique that cannot fully describe the complex nature of LNPs. Moreover, many methods rely on assuming LNPs adopt a spherical morphology, whereas recent work has suggested that LNPs likely adopt an elongated structure^15^.

Here, we analyze LNPs with an array of label-free solution-based biophysical techniques including sedimentation velocity analytical ultracentrifugation (SV-AUC), field-flow fractionation combined with multiangle light scattering (FFF-MALS), and size exclusion chromatography in-line with synchrotron small-angle X-ray scattering (SEC-SAXS). We apply these techniques in concert to characterize a library of gold standard LNPs that differ in their lipid excipients and formulation techniques. SV-AUC was implemented to determine the polydispersity of LNPs in terms of their mRNA loading and is used to contrast with the limited resolution of DLS. FFF-MALS provided more specific details on LNP physicochemical characteristics, such as hydrodynamic radius (R_h_), geometric radius (R_geometric_), lipid content, and mRNA loading, in a single run. SEC-SAXS was employed to identify particle shape, size, and key interactions and ordering within the LNP.

Finally, we demonstrate the utility of these strategies in predicting LNP transfection in primary human T cells, intravenous delivery, and intramuscular delivery, as well as discuss how the different formulation techniques impact the toxicity and endosomal escape properties of each LNP formulation. This work implements orthogonal techniques to uncover LNP characteristics and structure-activity relationships and establishes a workflow to probe LNPs with increasing resolution.

## Results

### Formulation and traditional characterization of the LNP library

Four representative LNP formulations were chosen based on their clinical significance and status as gold standards. Names were assigned according to the ionizable lipid, including MC3 (Alnylam), C12-200 (academic), SM-102 (Moderna), ALC-0315 (Pfizer/BioNTech), and were prepared with firefly (FLuc) luciferase mRNA. The MC3 formulation represents ONPATTRO®, the first siRNA-LNP therapy, utilized for hereditary transthyretin-mediated amyloidosis^16^. SM-102 and ALC-0315 correspond to the SPIKEVAX® and COMIRNATY® SARS-CoV-2 vaccines, respectively^17^. The C12-200 formulation is an academic gold standard for siRNA and mRNA delivery^18^.

Each LNP was formulated utilizing pipette mixing (*i.e.*, bulk mixing) or polydimethylsiloxane microfluidic devices. Microfluidic LNPs containing no mRNA were also created as controls. Encapsulated mRNA concentration and relative encapsulation efficiencies were determined by RiboGreen assay, where the microfluidic-formulated LNPs had mRNA concentrations of 40–60 ng/µL and encapsulation efficiencies >80%, while bulk mixed LNPs had lower mRNA concentrations of 10–30 ng/µL and encapsulation efficiencies of ∼50% (**Figure 1A–B**). One exception were the C12-200 LNPs, which maintained the same mRNA concentration of ∼45 ng/µL for both formulation methods. DLS in combination with static light scattering showed that microfluidic-formulated LNPs had ∼40 nm hydrodynamic radii with LNP concentrations of ∼10^12^ LNP/mL, whereas the bulk mixed LNPs had larger hydrodynamic radii of ∼100 nm and smaller LNP concentrations of ∼10^11^ particles/mL (**Figure 1C–D**). ζ-potential measurements indicated that all LNPs had a neutral charge (**Figure 1E**). Apparent p*K*_a_ was determined by a fluorescent 6-(*p*-toluidino)-2-naphthalenesulfonic acid (TNS) assay, and were ∼6 for all LNPs (**Figure 1F**). Cryogenic transmission electron microscopy (cryo-TEM) imaging was performed to evaluate LNP morphology (**Figure 1G**). Microfluidic-formulated LNPs maintained a uniform electron dense core with narrower size distributions than bulk mixed LNPs, which had more variable size and internal structure, with some aqueous compartments. Of note, the C12-200 LNP formulations had distinct multilamellar ring structures that were not observable for other lipid formulations that contained an amorphous core with an exterior bilayer.

**Figure 1.**
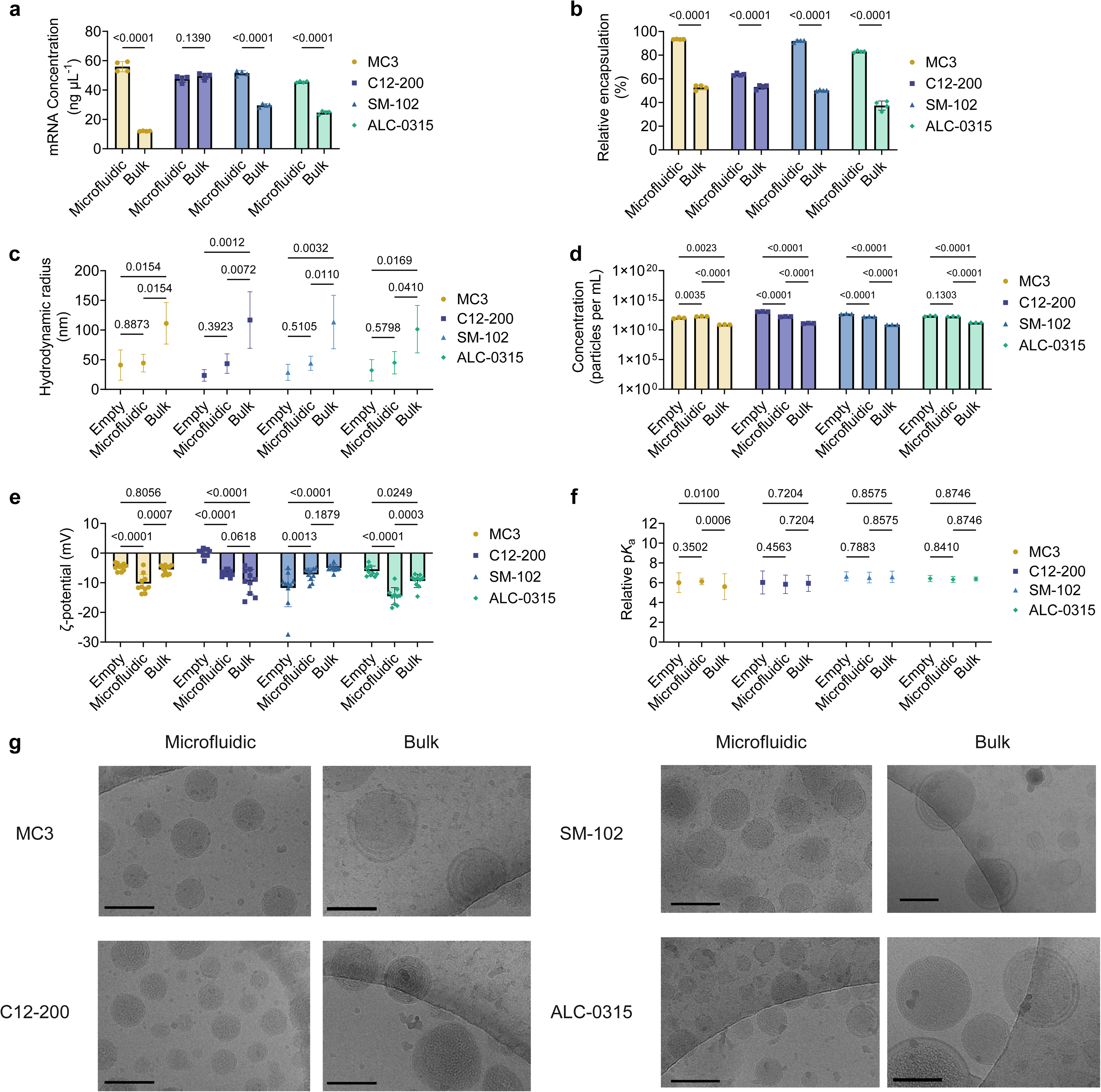
Traditional physicochemical characterization of LNPs. **A–G,** LNP physicochemical characteristics including, **(A)** mRNA concentration and **(B)** relative encapsulation efficiency determined by a RiboGreen assay, **(C)** hydrodynamic radius and **(D)** particle concentration obtained via DLS, **(E)** ζ-potential calculated via electrophoretic light scattering, **(F)** apparent p*K*_a_ determined by a TNS assay, and **(G)** morphology utilizing cryo-TEM. Measurements are reported as mean ± SD of *n* = 4 technical replicates for A–B, *n* = 3 technical replicates for C–D, and *n* = 10 for E. For F, SD was calculated from the upper and lower 95% confidence intervals (see Methods). Scale bar corresponds to 100 nm. **(A–F)** Two-way ANOVA with Holm–Šídák correction for multiple comparisons were used to compare the **(A)** concentration, **(B)** percent encapsulation, **(C)** radius, **(D)** concentration, **(E)** ζ-potential, and **(F)** p*K*_a_ across LNP formulations. Source data are provided as a Source Data file.

### Assessment of LNP polydispersity using analytical ultracentrifugation

SV-AUC is a quantitative, first principle approach that is sensitive to particle size, shape, and density^19^. Proper AUC implementation does not induce enough shear force to lyse the LNP and instead, can provide insights into LNP polydispersity^20^. Because this approach is sensitive to the density of the solutes examined, the tool can discriminate unloaded LNPs that float versus those loaded with the denser mRNA payload that increase sedimentation.

The results from this analysis are shown in **Figure 2A–D**. Each sample in the panel was characterized by DLS and subsequently analyzed by AUC. Modest rotor speeds were applied with the UV detection optimized for detection of ribonucleic acid bases (260 nm) at 20 °C. In the primary data, evidence for both flotation and sedimentation are readily observed (**Supplementary Figure 1**). A least-squares *g*(s)* model^21^ was applied to assess the presence of both floating and sedimenting species^22^. The multimodal appearance of the distributions suggests the existence of subpopulations in each LNP that were not detected by DLS. Most of the detected mass for the MC3, SM-102, and ALC-0315 LNPs were found to float, with S values spanning 0 to - 200S. In contrast, C12-200 LNPs predominantly sediment, with S values spanning 0 to ∼100S. These trends are in-line with the published densities of the MC3, C12-200, SM-102, and ALC-0315 ionizable lipids, 0.866, 0.948, 0.925, 0.919 g/cm^3^, respectively. The limited number of sedimenting species in MC3 is likely due to its lower density compared to the other ionizable lipids. In each case, the microfluidic samples display far less polydispersity than the bulk mixed samples.

**Figure 2.**
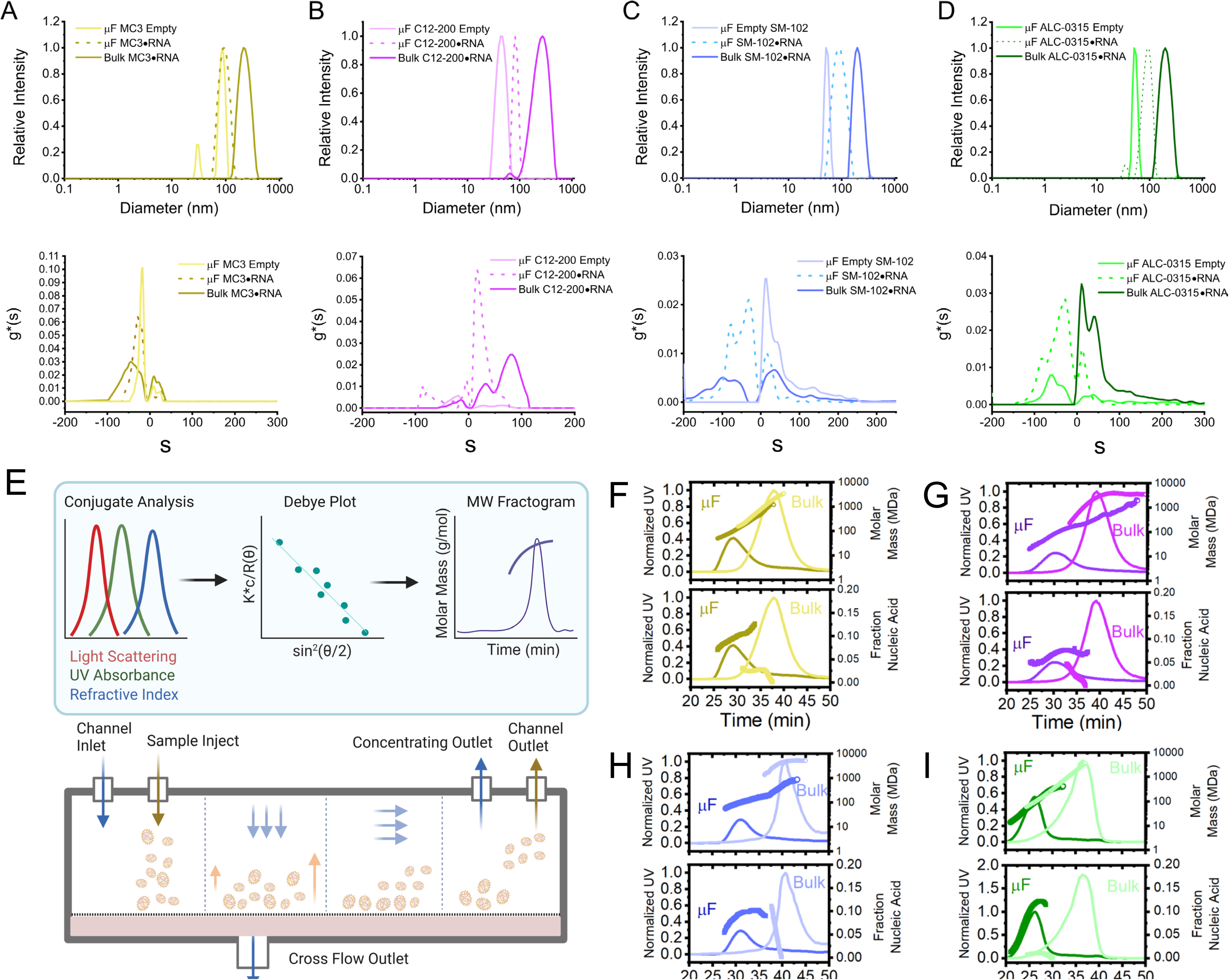
Analysis of LNPs by DLS, AUC, and FFF-MALS. **A–D,** In the upper panels, DLS distributions are shown for microfluidic empty LNPs (light solid line), microfluidic mRNA LNPs (dotted line), and bulk mixed mRNA LNPs (dark solid line) for each of the four ionizable lipids: **(A)** MC3, **(B)** C12-200, **(C)** SM-102, and **(D)** ALC-0315. Diameter in nanometers is shown on the x-axis and relative intensity on the y-axis. In the lower panels, ls-g*(s) distribution profiles from AUC analyses are shown for the same samples using the program SEDFIT. Species observed to the left of zero represent floating species while those to the right represent sedimenting species, with the area under the curve proportional to the relative abundance in the formulation. E, Diagram of FFF and LNP analysis (left). F–I, In each panel, both the determinedmolar mass profiles (upper panels) and mRNA weight fraction (lower panels) derived from MALS analysis for microfluidic (dark solid line) and bulk mixed (light solid line) LNPs were overlaid with UV fractograms from in-line FFF separations. Quadrants represent **(F)** MC3, **(G)** C12-200, **(H)** SM-102, and **(I)** ALC-0315. Additional parameters derived from these analyses are provided in Extended Data 1. **E,** Created in BioRender. Hamilton, A. (2024) https://BioRender.com/f01w633. Source data are provided in Zenodo (https://10.5281/zenodo.17042311).

### All-in-one size and compositional analysis using multiangle light scattering

MALS, coupled with UV and refractive index (RI) detection, provides physicochemical information independent of the effects of shape^23^. However, a key factor in its application is the separation method used prior to data collection, which is typically achieved via chromatography^24^. By applying FFF, superior separation of particles in the 1–1000 nm range, such as LNPs, is achieved under gentler conditions compared to chromatographic methods due to the absence of a stationary phase (**Figure 2E**)^25^.

The fractograms for each separation reflect the relative abundance of analytes fractionated by this method. In each case, a major peak representing the bulk of the LNP mass is observed during the separation. However, these profiles are relatively broad and, in some cases, asymmetric, indicating the presence of additional subpopulations in the formulation. The retention times for bulk mixed LNPs were broader and longer than those for LNPs prepared using microfluidics. As expected for polydisperse samples, the molar mass measured by FFF-MALS increased with retention time, spanning approximately 1–2 orders of magnitude in the megadalton range for all formulations examined, where bulk mixed samples displayed 10 to 20-fold higher masses (**Figure 2F–I**). mRNA and lipid mass concentrations were determined using a previously described strategy for correcting the concentration measurements for UV scattering from the LNPs^23^. Microfluidic LNPs contained 4–30 times higher mRNA concentration than bulk mixed LNPs. While the mRNA component is present throughout the microfluidic peak, in bulk samples it appears to partition into smaller subpopulations, disappearing rapidly with larger sizes.

Empty LNPs prepared by microfluidic methods were larger in mass and hydrodynamic radius (R_h_) compared to loaded microfluidic LNPs, with greater polydispersity as evidenced by M_w_/M_n_ (dispersity, Ð). Microfluidic LNPs displayed hydrodynamic radii (R_h_) that were 1.5 to 2-fold smaller than bulk mixed LNPs, and R_geometric_ that were 2–3-fold smaller. R_geometric_ provides the radius of a solid sphere that produces the same radius of gyration (R_g_), hence differing from the true R_g_ of an anisotropic particle, while R_h_ describes the effective radius of the particle based on its diffusion properties. Overall, FFF-MALS can better resolve the true dispersity of these polydisperse samples, as M_w_/M_n_ correlates more closely with the observed variations in mass and composition than the PDI values derived from DLS, which are based on intensity-weighted mean hydrodynamic diameters. A table of physicochemical parameters obtained by FFF-MALS is provided in **Extended Data Table 1**.

### Probing LNP internal structure and shape via small-angle X-ray scattering

SAXS is a free-solution measurement well-suited for probing the internal structure of LNPs because the scattering length density of the electron-dense nucleic acid component is significantly higher than lipids when exposed to X-rays. LNPs exhibit a characteristic first-order Bragg peak at a scattering vector (q) of approximately 0.1–0.15 Å⁻¹, corresponding to a distance (d) of ∼41.9–62.8 Å, where 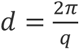, indicative of a highly ordered internal structure comprising tight interactions between RNA and lipid components. Prior work has established models for LNP structures composed of bilayers with a disordered core or multilamellar LNPs, which can be discerned by the spacing of the Bragg peaks^26–28^.

Most SAXS measurements of LNPs have been conducted under equilibrium conditions with high particle concentrations, without accounting for the polydispersity of the formulations. Since the scattering profiles represent the volume-weighted average of all scatterers within the X-ray path, we implemented two strategies in tandem to enhance the information content of the experiment: (i) in-line SEC and (ii) singular value decomposition (SVD) coupled with evolving factor analysis (EFA) to deconvolute the data (**Figure 3A**). SVD analysis allows for the identification of multiple significant species within the dataset, allowing for better representation of polydisperse mixtures. In comparison, static SAXS measurements are not amenable to this approach due to the lack of data density needed to resolve the components. After the principal components are identified, EFA is performed to estimate the specific frames or elution points of each species. Finally, the individualized SAXS profiles are extracted via regularized alternating least squares (REGALS) analysis, producing independent scattering data for each specie from the parent SEC peak^29^.

**Figure 3.**
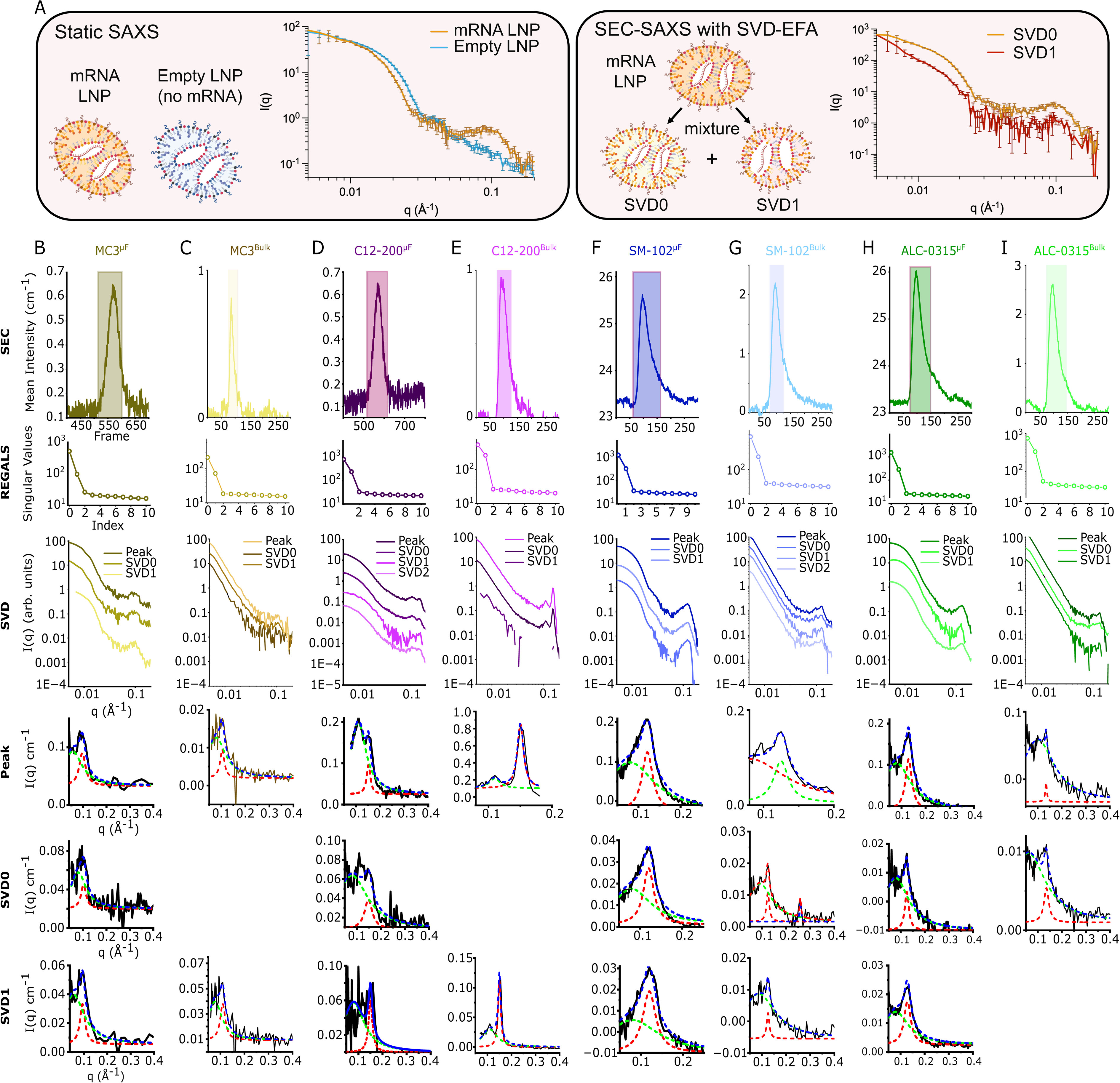
LNP batches contain discrete subspecies as identified by SEC-SAXS. **A,** Comparison between static SAXS and SEC-SAXS with single value decomposition (SVD) and evolving factor analysis (EFA). Measurements are reported as mean intensity ± SD, where SD is the rotational averaging of detector data. **B–I**, Individual SEC-SAXS parameters for **(B)** MC3^µF^, **(C)** MC3^Bulk^, **(D)** C12-200^µF^, **(E)** C12-200^Bulk^, **(F)** SM-102^µF^, **(G)** SM-102^Bulk^, **(H)** ALC-0315^µF^, **(I)** ALC-0315^Bulk^ LNPs. Each LNP underwent an identical processing protocol, involving identifying the main LNP peak in the SEC chromatogram (first row), selecting the number of singular values (second row), displaying scattering intensity of the main peak and corresponding singular values (third row), and fitting of the higher-q peak features (0.1 – 0.15 Å⁻¹) using multiple Lorentz peak feature (fourth, fifth, and sixth rows), where red is the first order Braggs peak fit, green represents higher-order particle disorder, and blue is the cumulative fit of the two features. Some SVD0 and SVD1 scattering plots were omitted due to low scattering signals. **A**, created in BioRender. Hamilton, A. (2025) https://BioRender.com/qx6vawp. Source data are provided in Zenodo (https://10.5281/zenodo.17042311)^50^.

For each LNP, we performed synchrotron SAXS measurements in-line with SEC, both with and without SVD analysis as incorporated in the program REGALS (**Supplementary Figure 2**). In control experiments, we obtained static measurements on microfluidic-formulated LNPs and did not observe concentration-dependent behavior or evidence of interparticle interference at LNP concentrations of ∼10¹¹ or lower (**Supplementary Figure 3**). Using UV-Vis data from an in-line diode array detector, raw data from this experiment showed strong UV absorbance at 260 nm, coinciding with X-ray scattering intensity between q ∼0.1–0.15 Å⁻¹ (**Supplementary Figure 4**). As a control, LNPs without mRNA were prepared and displayed no such peak feature. For each sample, 70–100 scattering profiles collected from the elution peak were combined for subsequent analysis, either by averaging the selected profiles of the isocratically eluted peak or by analyzing the profiles using SVD to decompose the data into minimal components with maximal redundancy (**Figure 3B– I, Supplementary Tables 1–4**). Two to three significant components were readily identified through SVD analysis for each LNP, and the data were decomposed accordingly. To substantiate the statistical uniqueness of the SAXS data, we utilized the Volatility of Ratio (V_r_) metric, which describes the variability in the ratio of scattering intensity between two profiles and is advantageous over other methods of profile comparison, especially in regimes of weaker signals^30^. Different degrees of low similarity were observed between experimental profiles with or without SVD analysis and by method of preparation, which indicates that our method results in statistically significant differences in the experimental profiles (**Figure 4A**).

**Figure 4.**
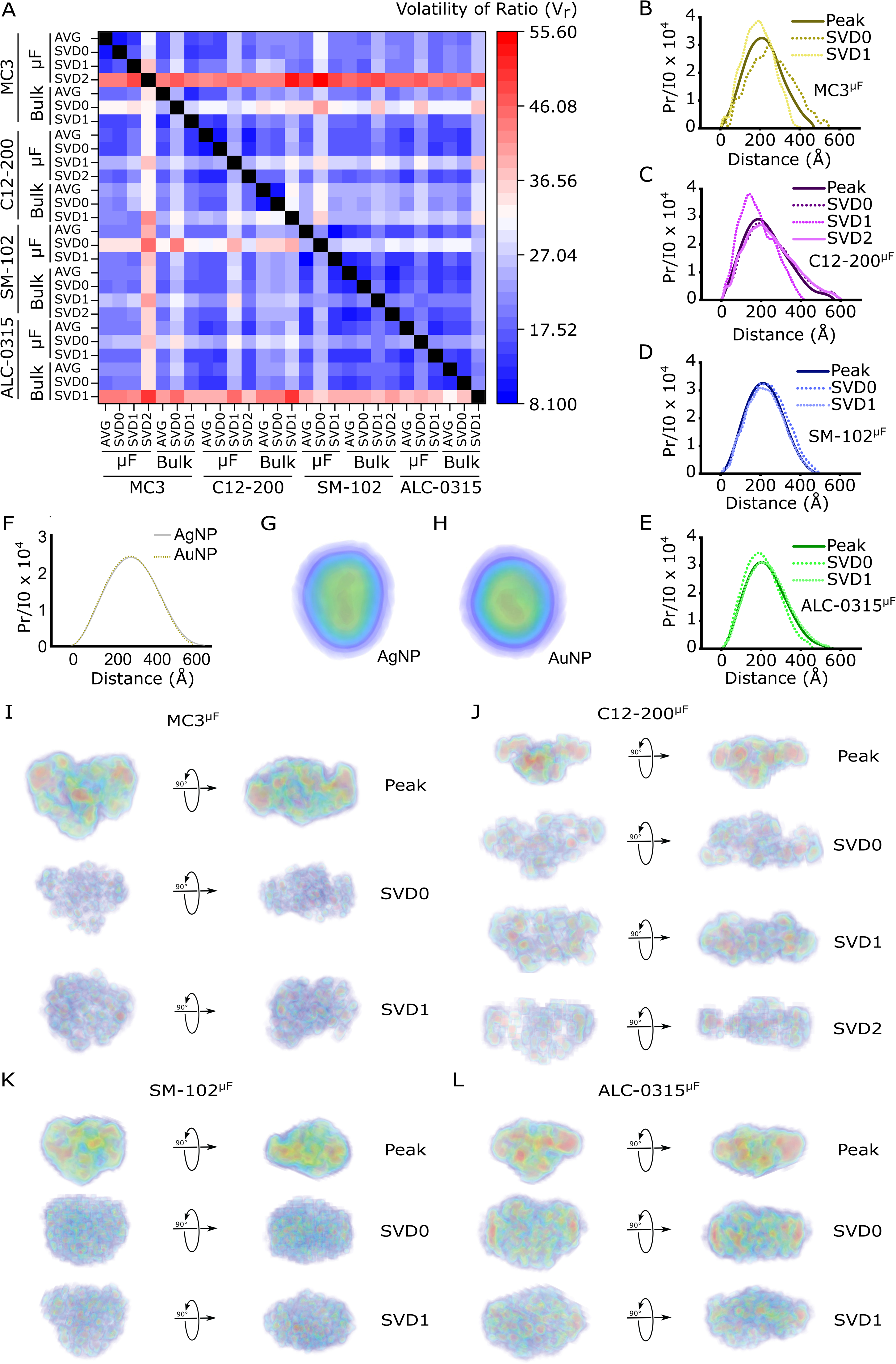
SEC-SAXS profiles have low degree similarity and show that LNPs adopt non-spherical morphologies. **A,** Volatility of Ratio (V_r_) between the SEC-SAXS average peak and individual SVDs. **B–E,** P(r) analysis of the peak and SVDs for **(B)** MC3, **(C)** C12-200, **(D)** SM-102, and **(E)** ALC-0315 microfluidic-formulated LNPs. **F,** P(r) analysis for 60 nm silver and gold spherical nanoparticles. **G–L,** DENSS *ab initio* electron density reconstructions from the SEC-SAXS profiles for **(G)** 60 nm silver spherical nanoparticles, **(H)** 60 nm gold spherical nanoparticles, **(I)** MC3^µF^ LNPs, **(J)** C12-200^µF^ LNPs, **(K)** SM-102^µF^ LNPs, and **(L)** ALC-0315^µF^ LNPs. P(r) analyses and DENSS reconstructions for bulk mixed LNPs could not be generated due to their larger size (𝑞_𝑚𝑖𝑛_ ∗ 𝐷_𝑚𝑎𝑥_ ≤ 𝜋). Source data are provided in Zenodo (https://10.5281/zenodo.17042311).

These data were further assessed in two complementary ways. First, the higher-q peak features between ∼0.1–0.15 Å⁻¹ were analyzed using multiple Lorentz peak fitting to deconvolve the signal of structural ordering from the overlapping disordered signal at larger length scales, providing measurements of peak width (FWHM) and magnitude (intensity at peak (I_p_), area)^6^. Additionally, the size and shape of the LNPs were evaluated using the P(r) pair distance distribution function. Although Guinier analysis was not possible, the accessible q-range (q_min_ = 0.005 Å⁻¹) allowed for the assessment of R_g_ and the maximum particle dimension (D_max_) up to ∼628 Å using P(r) analysis, in conditions where 𝑞_𝑚𝑖𝑛_ ∗ 𝐷_𝑚𝑎𝑥_ ≤ 𝜋 is satisfied (**Figure 4B–E**).

We found that the shape of the scattering profile and location of the peaks were primarily determined by the lipid type and not by the formulation method. Rather, the formulation method impacted the intensity of the peak, where higher intensity corresponds to a greater degree of lipid-RNA ordering. For each LNP, the first-order Bragg peak (q_peak_) is 0.10, 0.16, 0.13, and 0.13 Å^-1^, for MC3, C12-200, SM-102, and ALC-0315, respectively. The Bragg peak appeared only with RNA-loaded LNPs, demonstrating that the addition of the payload creates unique mesophases of 40–60 Å within the LNP. Of the four ionizable lipids examined in this study, C12-200 is the most dissimilar from the others due to its shorter alkyl tails and piperazine core, in contrast to only one nitrogen atom per molecule with longer alkyl tails for the other ionizable lipids. In fact, two peaks appear in the C12-200 q regime, where a smaller peak can be observed at q = 0.12 Å^-1^.

The multiple Lorentz peak fitting allowed for resolution of the primary peak feature from overlapping signal at lower q values. Using this approach, we observed that LNPs formulated with our microfluidic devices have higher intensity Bragg peaks for all LNPs except C12-200, upwards of ∼2-fold for the averaged scattering from the SEC peak and upwards of ∼10-fold higher for the SVD0 deconvoluted profiles; this higher intensity was concomitant with large peak area and width (**Supplementary Table 3**). The formulation method also impacted the peak position observed after the SEC enrichment step.

Shape distribution analysis was only possible for microfluidic-formulated LNPs as the bulk mixed LNPs did not satisfy the 𝑞_𝑚𝑖𝑛_ ∗ 𝐷_𝑚𝑎𝑥_ ≤ 𝜋 criterium. To assist in a qualitative assessment of these distributions, we also collected static SAXS measurements on 60 nm silver and gold nanoparticles (**Figure 4F–H, Supplementary Figure 5**). Relative to these isotropic particles, the shape distribution functions of LNPs displayed a greater skew of interatomic vectors to larger values relative to the peak (∼R_g_), suggestive of anisotropic shapes, with the greatest anisotropy observed for the C12-200 LNP. The algorithm DENsity from Solution Scattering (DENSS) can be applied to SAXS data from LNPs to reconstruct *ab initio* low resolution electron density in three dimensions^15^. Consistent with the trends observed by P(r) analysis, anisotropic ellipsoids are observed for all four microfluidic LNPs (**Figure 4I–L**). Utilizing the parameters from the P(r) analyses, we estimated a “shape factor” via calculating 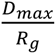, which equals ∼2.58 and ∼3.0 for spheres and prolate ellipsoids, respectively. By plotting the D_max_ vs R_g_ of each microfluidic LNP, we found that their values ranged from ∼2.9 for MC3 and SM-102 to 3.1 for C12-200 and ALC-0315, consistent with prolate ellipsoidal morphology (**Supplementary Table 5, Supplementary Figure 6**). Moreover, for C12-200, SM-102, and ALC-0315, there were differences in the shape factor between the average, SVD0, and SVD1 profiles, indicating that some LNP subpopulations have polydispersity in their shape.

To explore how diverse types of cargo impact LNP structure, we encapsulated green fluorescent protein (GFP) siRNA using the MC3 formulation to mimic ONPATTRO®. We evaluated microfluidic and bulk mixed MC3 siRNA LNPs first using DLS (**Supplementary Figure 7**). Utilizing our SEC-SAXS methodology, we then analyzed the LNPs, where after SVD, MC3-siRNA^µF^ had one significant specie (**Supplementary Figure 8A–C**) whereas MC3-siRNA^Bulk^ had two species (**Supplementary Figure 8D–F)**. Unlike the mRNA LNPs, MC3-siRNA^Bulk^ had a more intense q_peak_ than MC3-siRNA^µF^, meaning LNP internal structure is sensitive to cargo type and that formulation methods for a given LNP are not consistent in creating similar RNA-lipid interactions when switching RNAs. (**Supplementary Figure 8G–I**). The DENSS reconstructions for MC3-siRNA^µF^, showed an elongated spherical morphology, similar to the mRNA MC3 LNPs (**Supplementary Figure 8J**).

### Evaluation and correlation in biological models

Current efforts to formulate next-generation LNPs rely on high-throughput screening of hundreds or thousands of unique LNPs to identify promising candidates for the desired biological application^31,32^. Due to a lack of structure-activity relationships between the lipid components and mRNA translation, lipid and LNP design are often randomly explored leading to a substantial percentage of the tested library to perform poorly. Attempts to correlate biological performance with LNP physicochemical characteristics and structure have yielded unsatisfactory associations because of the utilization of low-resolution analytical techniques. Thus, we determined whether our solution-based biophysical techniques could reveal stronger correlations between LNP characteristics and efficacy. We evaluated our lipid library in primary human T cells, intravenous (IV) administration, and intramuscular (IM) administration. CD4+ and CD8+ T cells (1:1) were obtained from healthy human donors and activated for 24 h using CD3/CD28 human Dynabeads^TM^. The eight LNPs encapsulating FLuc mRNA were incubated with the activated T cells, and after 24 h, luminescence and viability were analyzed (**Figure 5A–B**). While the MC3^µF^, SM-102^µF^, and ALC-0315^µF^ LNPs had 2–3-fold higher luminescent signal than their bulk mixed counterparts, the C12-200^Bulk^ LNP outperformed the C12-200^µF^ LNP by almost 3-fold and was the highest performing LNP out of the library. Additionally, the bulk mixed LNPs had 10–20% higher viability than the microfluidic-formulated versions except for the C12-200 group, which had similar viabilities of ∼80%.

**Figure 5.**
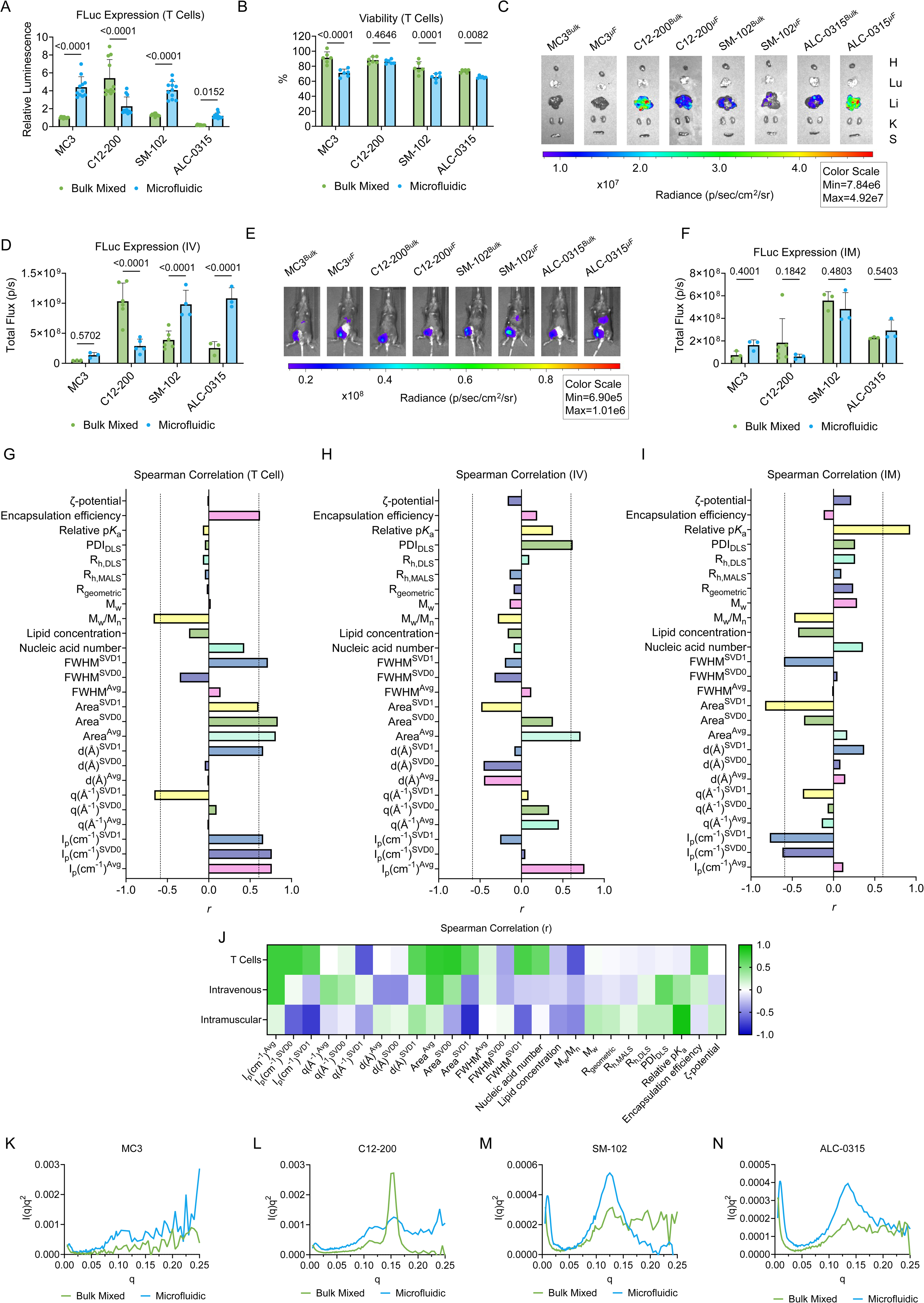
LNP transfection in biological models and correlation with physicochemical parameters. **A–B,** MC3, C12-200, SM-102, and ALC-0315 LNPs, either bulk mixed or microfluidic-formulated with firefly luciferase mRNA, were incubated in 1:1 CD4+:CD8+ human primary T cells obtained from healthy donors at a dose of 300 ng per 60,000 cells. After 24 h, **(A)** luminescence and **(B)** viability were quantified. Relative luminescence and viability are reported as mean ± SD of *n* = 11 for luminescence and *n* = 6 for viability. **C–D,** The eight LNPs were administered intravenously into C57BL/6J mice at a dose of 0.1 mg of mRNA per kg of body mass (mpk). After 6 h, mice were dissected, and organ luminescence was **(C)** imaged and **(D)** quantified using an in vivo imaging system (IVIS). Heart (H), lungs (Lu), liver (Li), kidneys (K), and spleen (S), were dissected and analyzed. Total flux is reported as mean ± SD of *n* = 6 for the C12-200^Bulk^ and SM-102^Bulk^, *n* = 5 for the C12-200^µF^, *n* = 4 for SM-102^µF^, and *n* = 3 for the MC3 and ALC-0315 groups. **E–F,** The eight LNPs were also administered intramuscularly into C57BL/6J mice at a dose of 0.05 mpk. After 6 h, luminescence at the injection site was **(E)** imaged and **(F)** quantified using IVIS. Total flux is reported as mean ± SD of *n* = 6 for the C12-200^Bulk^ and *n* = 3 for all other groups. **G–J,** Spearman correlations for the **(G)** T cell, **(H)** intravenous, and **(I)** intramuscular luminescence values utilizing the physicochemical parameters from the traditional, FFF-MALS, and SEC-SAXS methods, with **(J)** a heat map representing the entire data set. For the Spearman correlation graphs, dotted lines represent *r* = −0.6 and 0.6. **K–N**, Overlayed Kratky plots of microfluidic and bulk mixed **(K)** MC3, **(L)** C12-200, **(M)** SM-102, **(N)** ALC-0315 LNPs obtained via SEC-SAXS. **A–B,D,F,** Two-way ANOVA with *post hoc* Holm–Šídák correction for multiple comparisons were used to compare bulk mixed against microfluidic-formulated luminescent values for each LNP group. Source data are provided as a Source Data file for **A–B,D,F**. Source data are provided in Zenodo (https://10.5281/zenodo.17042311) for **K–N**.

Upon injecting the FLuc LNPs intravenously into C57BL/6 mice and analyzing liver luminescence after 6 h, the transfection trends mirrored the T cell results (**Figure 5C–D**). The MC3^µF^, SM-102^µF^, and ALC-0315^µF^ LNPs induced 2–3-fold higher liver luminescence signal than the bulk mixed LNPs, whereas the C12-200^Bulk^ LNP outperformed C12-200^µF^ LNP by 3-fold. In contrast, intramuscular administration resulted in much closer luminescence at the injection site between the microfluidic and bulk mixed LNPs, with no statistically significant differences between the groups (**Figure 5E–F**).

Physicochemical information of the LNP library were correlated with the luminescence readouts. Utilizing Spearman correlations, with 𝑟 > 0.6 or 𝑟 < −0.6 as a threshold for medium-to-strong correlation, several physicochemical factors emerged as having strong correlation with biological outcome (**Figure 5G–J; Supplementary Table 6**). T cell and IV luminescence had strong positive correlation with the intensity of the average SEC-SAXS peak (I_p_ ^Avg^), with both the SVD0 (I_p_ ^SVD0^) and SVD1 (I_p_ ^SVD1^) peaks also correlating with the T cell data but not the IV data. When comparing LNPs based on formulation method, the I_p_ is predictive of LNP performance, as the formulation method with the stronger peak induced greater luminescence in T cells and IV delivery. This may explain the counterintuitive C12-200 results, as C12-200^BM^ elicited a stronger peak than C12-200^µF^, while the opposite occurred for the other three LNP groups (**Figure 5K–N**). In contrast, I_p_ ^SVD0^ and I_p_ ^SVD1^ produced a strong negative correlation with the IM data, indicating an inverse relationship. Moreover, the T cell and IV biological results had a strong positive correlation with the area of the SAXS peak, particularly for the average peak (Area^Avg^), with T cells also showing a significant relationship with the SVD0 peak area (Area^SVD0^) and SVD1 peak area (Area^SVD1^). Between the biological models, the T cell, but not IV results, also correlated with the position of the SAXS peak maximum at 0.1–0.2 q for SVD1 (q^SVD1^), with lower q indicating more efficacious delivery.

Encapsulation efficiency and nucleic acid number were determinant factors for T cells, but not the *in vivo* models. Additionally, p*K*_a_ has a strong correlation solely for IM delivery. R_h_ and R_geometric_ were poor predictors of performance for all three biological models, and instead, polydispersity from MALS was a more accurate measure, with lower M_w_/M_n_ correlating with stronger luminescence than size. Counterintuitively, for IV delivery, PDI has a strong positive correlation, indicating inconsistency between the MALS and DLS measurements.

We also investigated LNPs in cancer cells, which have a distinctive cellular biology, especially their cellular metabolism and proliferation^33^. Utilizing FaDu cells, an oral squamous cell carcinoma line, we incubated the eight LNPs at a dose of 20 ng per 20,000 cells. After 24 h, we analyzed the luminescence and found that the LNP trends were unique compared to the studies with T cells (**Supplementary Figure 9A**). MC3^Bulk^, SM-102^Bulk^, and ALC-0315^Bulk^ outperformed the corresponding microfluidic versions, whereas C12-200^Bulk^ and C12-200^µF^ had similar luminescence values. None of the LNPs induced less than 75% viability (**Supplementary Figure 9B**). When correlating transfection with physicochemical parameters, we found that larger size and positive ζ-potential correlated with increased luminescence, whereas high relative encapsulation efficiency, larger SAXS peak intensity, and wider area negatively correlated with luminescence (**Supplementary Figure 9C**). These differences are likely due to the higher rates of endocytosis in cancer cells, which may allow larger LNPs, such as the bulk mixed formulations, to more easily transfect these cells^34,35^.

### Assessing *in vivo* LNP toxicity

Measuring toxicity is equally important in studying LNP compositions and formulation methods. To do this, we reformulated the LNPs with 5moU erythropoietin (epo) mRNA. Epo is a glycoprotein hormone that stimulates red blood cell production, and has been utilized to assess LNP-induced toxicity^36^. All eight formulations showed similar but not exact R_h_ and relative encapsulation compared to the same LNPs with FLuc mRNA (**Supplementary Figure 10**). Mice were then intravenously administered the LNPs at a dose of 0.4 mpk, four times higher than the FLuc mRNA dose from the previous study. While no complications were observed with the bulked mixed and microfluidic FDA-approved formulations, both C12-200 formulations induced severe lethargy in all mice, with two out of the six mice dying in each group (**Figure 6A–B**).

**Figure 6.**
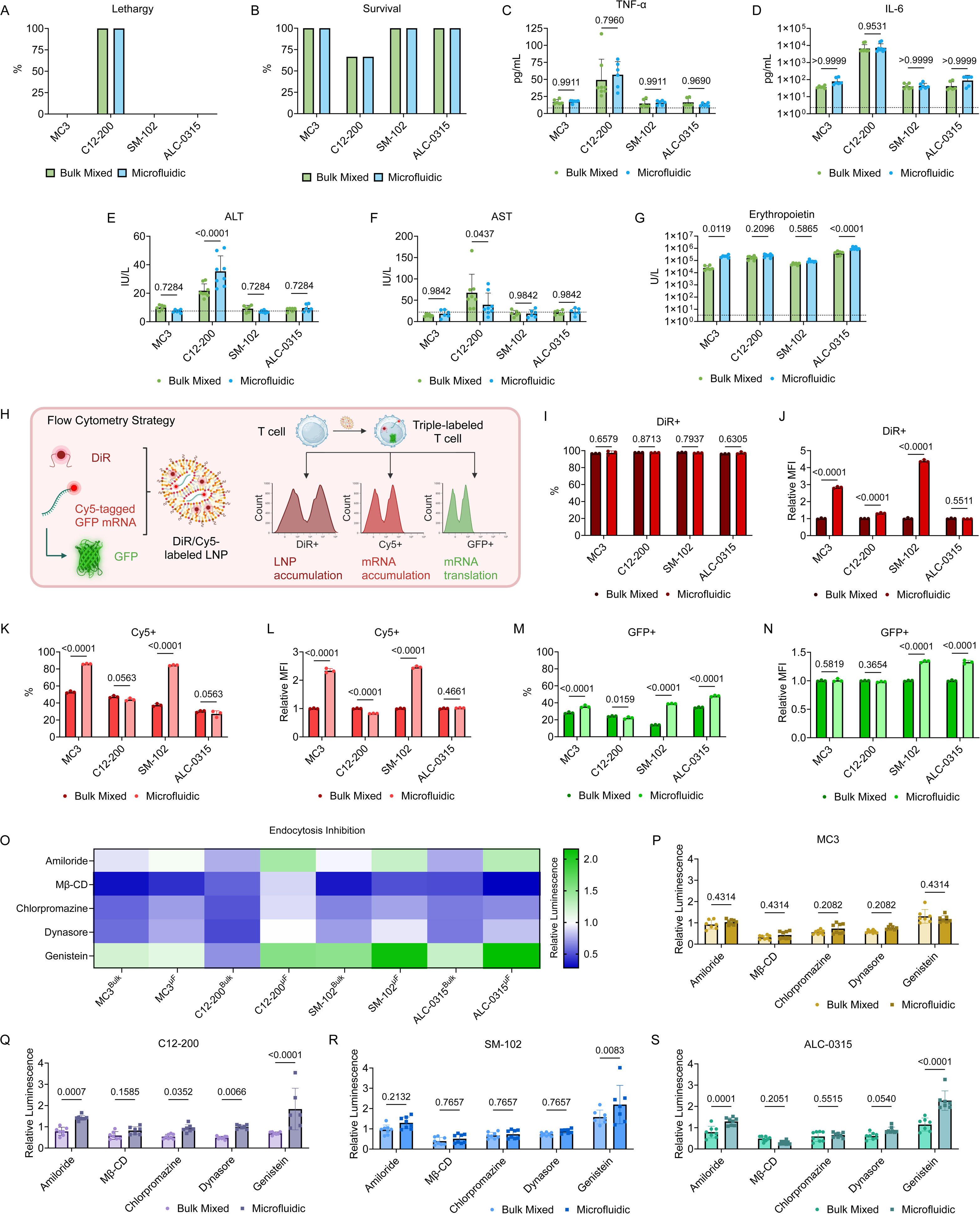
Analysis of LNP toxicity and endosomal escape. **A–G,** C57BL/6J mice were intravenously administered with the eight LNPs encapsulating epo mRNA at 0.4 mpk. Mice were monitored for 1 h for **(A)** indications of lethargy and **(B)** survival. After 6 h, blood was drawn, serum was isolated, and **(C)** TNF-α, **(D)** IL-6, **(E)** ALT, **(F)** AST, and **(G)** epo levels were ascertained by ELISAs. Survival and lethargy are reported as percentage of total of *n* = 6 for the C12-200 groups and *n* = 3 for all other groups. Serum proteins levels are reported as mean ± SD of *n* = 8 for the C12-200 groups, except for TNF-α for C12-200^µF^ which is *n* = 6, and *n* = 6 for all other groups. **H,** All eight LNPs were reformulated with 0.5 mol% DiR and Cy5-tagged GFP to examine LNP uptake, mRNA uptake, and mRNA translation in T cells. **I–N,** Human primary T cells (1:1, CD4+/CD8+) were activated for 24 h with human CD3/CD28 Dynabeads and then incubated with the dual-fluorescent GFP LNPs at a dose of 600 ng per 250,000 cells. After 24 h, the T cells were analyzed by flow cytometry to measure **(I)** %DiR positive cells, **(J)** DiR MFI, **(K)** %Cy5 positive cells, **(L)** Cy5 MFI, **(M)** %GFP positive cells, and **(N)** GFP MFI. **O–S,** Activated T cells were incubated with endocytosis inhibitors for 30 min. After washing, the cells were dosed with the eight LNPs reformulated with FLuc mRNA at a concentration of 300 ng per 60,000 cells. After 24 h, **(O)** luminescence was quantified relative to treated T cells with no inhibitors, and comparisons were plotted for the **(P)** MC3, **(Q)** C12-200, **(R)** SM-102, and **(S)** ALC-0315 LNPs. Relative luminescence is reported as mean ± SD of *n* = 7 or 8. **C–G,I–N,P–S,** Two-sided multiple unpaired t test with *post hoc* Holm–Šídák correction for multiple comparisons were used to compare bulk mixed against microfluidic-formulated values for each LNP group. **H** created in BioRender. Hamilton, A. (2025) https://BioRender.com/2a8cgih. Source data are provided as a Source Data file.

Additionally, after 6 h, serum was isolated from each mouse to examine the inflammatory cytokines TNF-α and IL-6, liver toxicity markers alanine transaminase (ALT) and aspartate transaminase (AST), and epo levels. Consistent with the observed toxicity, the FDA-approved formulations induced only 2-fold increases in TNF-α levels compared to control mice while the levels for C12-200 increased by 7–8-fold, with no statistical differences between C12-200^Bulk^ and C12-200^µF^ (**Figure 6C**). All LNPs increased IL-6 levels with the FDA-approved LNPs increasing IL-6 by 10–25-fold, compared to the C12-200 LNPs that increased levels by 1,000-fold (**Figure 6D**). Similar to TNF-α, no major differences in IL-6 concentration were observed between formulation methods. The ALT and AST results differed slightly from this trend. While both C12-200 LNPs induced higher levels of AST and ALT, the C12-200^Bulk^ AST levels were 2-fold greater than C12-200^µF^, whereas the opposite trend was observed for ALT levels (**Figure 6E–F**).

The toxicity is likely due to the structures of the ionizable lipids as MC3, SM-102, and ALC-0315 all contain biodegradable ester groups whereas C12-200 has no biodegradable moieties, which are essential for mitigating the harmful effects of the lipids^37^. The increased levels of cytokines and liver markers were not a result of greater hepatic epo production, as ALC-0315^Bulk^ induced a similar amount of epo as the C12-200 LNPs and ALC-0315^µF^ was 10-fold greater (**Figure 6G**). Secondly, ALC-0315^Bulk^ produced 10-fold less epo than ALC-0315^µF^, yet maintained the same TNF-α, IL-6, AST, and ALT values as ALC-0315^µF^. Among the formulation methods, MC3^µF^ also outperformed MC3^Bulk^, whereas there were no statistical differences between the C12-200 and SM-102 formulations. The different trends for the LNPs with epo mRNA and the LNPs with FLuc mRNA are likely based on the disparity in mRNA lengths, being 859 and 1,922 nucleotides (nt), respectively. We have previously found that LNP trends are different depending on whether the LNPs encapsulate FLuc or epo mRNA^38^. Moreover, since epo was measured in the serum, there may be LNP-induced effects that alter the biochemical pathways that govern epo secretion into the bloodstream.

### Examining the role of endosomal escape

Finally, we investigated the relationship between endosomal escape and the different LNP formulations. For most applications of LNPs, endosomal escape is the final barrier for delivery. It is estimated that only a fraction of particles in endosomes can translocate their cargo to the cytosol, which is crucial as mRNA will rapidly degrade during lysosomal maturation^39^. Utilizing T cells as our model, we reformulated all eight LNPs with Cy5-tagged 5moU GFP mRNA and the lipophilic near-infrared carbocyanine dye, DiR, at 0.5 mol%. This enabled the LNPs to be monitored in three simultaneous ways, with DiR, Cy5, and GFP measuring LNP uptake, mRNA uptake, and mRNA translation, respectively (**Figure 6H, Supplementary Figure 11**). We incubated the LNPs into T cells at 600 ng per 250,000 cells and analyzed fluorescence by flow cytometry after 24 h. Despite all formulations successfully delivering DiR to over 95% of cells, the Cy5 positive cells ranged from 30% to 80% depending on the LNP (**Figure 6I–L**). This discrepancy could be due to large populations of DiR-containing LNPs that do not contain mRNA. When re-dosed at 50 ng per 250,000 cells, all eight formulations induced at least 80% DiR positive cells with Cy5+ cells ranging from <5% to 20% and no GFP positive populations greater than 7% (**Supplementary Figure 12A–C**). Despite the high proportion of DiR positive cells, differences in LNP uptake were observed by analyzing the median fluorescent intensity (MFI). As DiR was loaded as a mol% based on the ionizable lipid, different amounts of DiR were added to each of the four recipes, and thus MFI was only compared between preparation methods for each LNP. We found that MC3^µF^, C12-200^µF^, and SM-102^µF^ had 1.5–5-fold higher MFI than their bulk mixed counterparts, whereas the MFI for the ALC-0315 LNPs was approximately equal. A similar result was observed for Cy5 in both the percentage of cells and the MFI, with MC3^µF^ and SM-102^µF^ having 40% more positive cells and double the MFI compared to MC3^Bulk^ and SM-102^Bulk^, and the ALC-0315 LNPs having ∼30% Cy5 positive cells and equal MFI. However, although C12-200^µF^ facilitated more DiR uptake, it delivered 10% less mRNA than C12-200^Bulk^. Thus, while C12-200^µF^ may be better at facilitating LNP uptake, C12-200^Bulk^ is superior at delivering mRNA into the cell.

For GFP expression, the results aligned similarly with the FLuc luminescence, with C12-200^Bulk^ outperforming C12-200^µF^, whereas for the other formulations, the microfluidic version transfected more cells than the bulk mixed LNPs (**Figure 6M–N**). In comparing the GFP and FLuc studies, ALC-0315^µF^ was the top performing LNP for GFP expression but the worst microfluidic LNP for FLuc delivery. The 996 nt GFP mRNA has a similar length to the 859 nt epo mRNA and in both cases, the ALC-0315^µF^ was the top candidate, although for the other LNPs there were differences in the relative epo and GFP performances. ALC-0315^µF^ is a unique case in that the Cy5 uptake was similar to ALC-0315^Bulk^, but the latter LNP had 20% less GFP transfection. Enhanced endosomal escape is the likely mechanism for the discrepancies in ALC-0315 performance, whereas for the other LNPs, a combination of uptake pathways and endosomal escape are driving factors. The MC3 and SM-102 are examples of the dual importance of uptake and endosomal escape, as MC3^µF^ and SM-102^µF^ induced ∼80% Cy5 positive cells, whereas only 30–40% of these cells were GFP positive. We then compared the Cy5% and GFP% of the microfluidic LNPs with their shape factor. Comparisons of the bulk mixed LNPs were not possible as their shape factor could not be calculated due to their large sizes that did not satisfy the 𝑞_𝑚𝑖𝑛_ ∗ 𝐷_𝑚𝑎𝑥_ ≤ 𝜋 criteria. We found that as the shape factor increased, indicating elongation of the particle, mRNA uptake decreased sharply; however, for GFP translation, no such trend was observed (**Supplementary Figure 13A–B)**.

To further examine the role of endocytosis, we performed an endosome inhibitor study. T cells were activated and then incubated with compounds that inhibit a certain endocytosis pathway, including amiloride (macropinocytosis), methyl-β-cyclodextrin (lipid-raft-mediated endocytosis), chlorpromazine (clathrin-mediated endocytosis), Dynasore (dynamin-dependent endocytosis), and genistein (caveolae-mediated endocytosis).

Inhibitors were incubated for 30 min and then dosed with the FLuc LNPs. After 24 h, luciferase was measured, and luminescence was normalized for each individual formulation to treated control wells that did not receive inhibitors (**Figure 6O–S**). Methyl-β-cyclodextrin treatment resulted in sharp decreases in luminescence for all LNPs, but most strongly for the MC3 formulations, SM-102^Bulk^, and ALC-0315^µF^, suggesting that lipid-raft endocytosis is a major pathway for successful mRNA translation in T cells. No decreases in viability were observed for T cells treated with inhibitors, indicating that decreases in viability are a result of changes in endocytosis (**Supplementary Figure 14**). Chlorpromazine and Dynasore mostly inhibited luciferase expression, with the bulk mixed formulations more strongly impacted.

For amiloride, the bulk mixed LNPs had lower luminescence while the microfluidic LNPs had higher luminescence, indicating that formulation method can alter endosomal escape, although the magnitude of these differences depended on the formulation. Increases in luminescence are likely due to some pathways being dead ends for translation, where the LNPs are unable to escape the endosome. By removing these pathways, LNPs are more likely to transverse endocytosis pathways that lead to endosomal escape and therefore mRNA translation. Genistein increased luminescence for all LNPs except for C12-200^Bulk^, the latter having a 2-fold increase in luminescence upon treatment with genistein. This suggests that C12-200^Bulk^ can undergo escape in caveolae-mediated endosomes, allowing for increased transfection, which may also explain its enhanced delivery compared to C12-200^µF^. While the SM-102 and ALC-0135 formulations all had higher luminescence after genistein treatment, SM-102^µF^ and ALC-0315^µF^ demonstrated significantly higher luminescence compared to the bulk mixed counterparts.

## Discussion

Our results demonstrate that for formulations encapsulating the same mRNA cargo, different spatial properties are observed, indicating that lipid composition and formulation method are strong determinants of particle properties. Shape could be a key parameter for the interaction of mRNA-loaded LNPs with their target cells. A higher surface-to-volume (S/V) ratio increases the surface area available for interactions with the cell membrane, thereby enhancing the likelihood of efficient endocytosis. Smaller LNPs with high S/V ratios would be expected to diffuse more rapidly through the extracellular matrix, reducing resistance to movement and enabling them to reach the target cells more efficiently.

Along with overall shape, the internal structure of the lipid-RNA interaction is an important factor for RNA translation in T cells and for intravenous administration, as demonstrated by our Spearman correlations. More broadly, our correlation analyses point to the role of the biological environment in determining which physicochemical parameters are most essential for efficacious mRNA delivery. LNPs that enter the bloodstream are coated in a more complex protein corona than in media, which directly impacts both the uptake pathway and endosomal escape. This may explain the poor correlation between *in vitro* and *in vivo* LNP transfection^40^.

When comparing the microfluidic and bulk mixing techniques, clear differences emerge, despite utilizing the same excipients and molar ratios in each formulation. For example, in T cells, all FDA-approved LNPs have enhanced mRNA transfection when formulated using microfluidics, whereas C12-200^Bulk^ achieved superior delivery compared to C12-200^µF^. This suggests that the structure of the lipid components may be an important parameter in choosing the formulation method. It is likely feasible to optimize device composition, geometry, and flow to achieve specific LNP physicochemical properties. Since microfluidic formulation offers a potential path towards global scale up of mRNA LNP therapeutics, as has been demonstrated with our devices, effort should be placed on how physicochemical characteristics are altered on both small and large scales^41^.

While some predictive efforts utilize artificial intelligence (AI) to design ionizable lipids, our studies indicate that two identical recipes, formulated differently, can have strong differences in biological activity^42^. We envision that a combination of lipid structure-activity relationships combined with deep understanding of how formulation techniques relate to LNP properties is the key to unlocking rational LNP design. As such, we recommend the following insights to help guide LNP design and provide additional areas that should be investigated to help advance future LNP platforms:

- Formulation method – we demonstrate the importance in testing multiple formulation methods for a given LNP to optimize its performance. It is common to screen large libraries using bulk mixing and then switch to microfluidic devices for the top performers, but this may lead to false negatives.
- Shape – our results suggest that different formulations have unique shapes and this should be further investigated as it likely impacts protein corona formation^43^, cellular uptake^44^, and endocytosis^45^.
- Loading – our studies support a growing body of evidence that a large fraction of LNPs have no mRNA. Designing formulation methods and lipids that can encapsulate mRNA more uniformly would enhance the likelihood of successful mRNA delivery.
- RNA-lipid interaction – we show that having a higher degree of RNA-lipid ordering can enhance delivery in certain applications, including in primary T cells.
- Biological environment – by comparing distinctive biological models such as T cells and cancer cells, we showcase that the route of delivery and biological environment strongly impact successful mRNA delivery. Moreover, certain therapies, like vaccines, involve complex cellular and biochemical pathways that can effect LNP efficacy^46^.
- Endosomal escape – since previous studies have indicated that different cells have unique endosomes and endocytosis pathways, it is important to investigate the endocytosis mechanisms for the biological target of interest and optimize the lipids for the specific biological target^47,48^.
- Cargo – we have found that the relative performance of a given LNP formulation changes with different mRNA cargo. Moreover, it is known that the performance of LNPs is altered depending on the mRNA modifications and purity^49^. Therefore, it may be important to screen the desired mRNA of interest, rather than relying on initial screening with gene reporters.

Although the biophysical techniques implemented in this manuscript provide enhanced resolution compared to traditional techniques, they are limited in their cost and availability. For SV-AUC, standard ultracentrifugation in density gradient tubes followed by downstream UV/vis and fluorescent analysis would provide similar quality data and allow for the isolation of LNP fractions. MALS has gained popularity over the past decades and thus these instruments are more common, and are built in with in-line FFF or SEC systems. Obtaining high magnitude synchrotron radiation is only possible in select institutions around the world; still, several companies have developed benchtop SAXS instruments that provide quality scattering data. Some systems include options for in-line attachment of separating columns and 96-well plate formats for high-throughput screening.

In summary, we present a set of solution-based biophysical techniques that can analyze LNP polydispersity, composition, size, and shape with higher resolution. When paired with biological data, these analytical tools offer the ability to create rule sets that may guide future LNP design by establishing structure-activity relationships based on physicochemical properties.

## Acknowledgments

K.G. acknowledges support from the Johnson Research Foundation. The SV-AUC experiments were performed at the Johnson Foundation Biophysical and Structural Biology Core Facility (University of Pennsylvania, Philadelphia PA). The LiX beamline is part of the Center for BioMolecular Structure (CBMS), which is primarily supported by the DOE Office of Biological and Environmental Research (KP1605010). As part of NSLS-II, a national user facility at Brookhaven National Laboratory, work performed at the CBMS is supported in part by the U.S. Department of Energy, Office of Science, Office of Basic Energy Sciences Program under contract number DE-SC0012704. Additionally, the authors thank Emily Cento, Zhilin Chen, Max A. Eldabbas, and Emileigh Maddox of the Human Immunology Core and the Division of Transfusion Medicine and Therapeutic Pathology at the Perelman School of Medicine at the University of Pennsylvania for providing de-identified CD4+ and CD8+ T cells that were purified from healthy donor apheresis using StemCell RosetteSep™ kits. Cryo-EM imaging was provided by the Beckman Center for Cryo Electron Microscopy at the University of Pennsylvania Perelman School of Medicine. M.J.M. acknowledges support from a Burroughs Wellcome Fund Career Award at the Scientific Interface (CASI), an American Cancer Society Research Scholar Grant (RSG-22-122-01-ET) and a US National Science Foundation CAREER Award (CBET-2145491).

## Author Contributions

M.S.P., S.J.S., M.K., X.Z., M.C., J.B., and K.G. conceived and designed the experiments. M.S.P., S.J.S., A.H., M.K., X.Z., J.B., H.M.Y., A.R., R.A.J., and K.M. performed the experiments. M.S.P., S.J.S., M.K., X.Z., M.C., J.B., and K.G. analyzed the data. M.S.P. and K.G. wrote the manuscript. D.I. and M.J.M. edited the manuscript. All authors discussed the results and commented on the manuscript.

## Methods

All animal use was in accordance with the guidelines and approval from the University of Pennsylvania’s Institutional Animal Care and Use Committee (IACUC; protocol #806540).

### Materials

Lipid excipients were obtained from Avanti Polar Lipids (Alabaster, AL, USA). Firefly luciferase mRNA and erythropoietin mRNA was purchased from TriLink Biotechnologies (San Diego, CA, USA) and contained 5-methoxyuridine substitutions. Cy5-tagged green fluorescent protein (GFP) mRNA was purchased from APExBIO (Houston, TX, USA). GFP siRNA was purchased from Dharmacon (Lafayette, CO, USA). Dialysis cassettes, pH 7.4 10X PBS, and 1,1’-dioctadecyl-3,3,3’,3’-tetramethylindotricarbocyanine iodide (DiR) were obtained from Thermo Fisher Scientific (Waltham, MA, USA). Citrate buffer was purchased from Teknova (Hollister, CA, USA). Syringe filters were obtained from Genesee Scientific (El Cajon, CA, USA).

### Lipid nanoparticle preparation

Lipid excipients were dissolved in ethanol, utilizing the following molar ratios:

MC3 LNPs: D-Lin-MC3-DMA (50 mol%), DSPC (10 mol%), cholesterol (38.5 mol%), and DMG-PEG2000 (1.5 mol%).

C12-200 LNPs: C12-200 (35 mol%), DOPE (16 mol%), cholesterol (46.5 mol%), and C14-PEG2000 (2.5 mol%).

SM-102 LNPs: SM-102 (50 mol%), DSPC (10 mol%), cholesterol (38.5 mol%), and DMG-PEG2000 (1.5 mol%).

ALC-0315 LNPs: ALC-0315 (46.3 mol%), DSPC (9.4 mol%), cholesterol (42.7 mol%), and ALC-0159 (1.6%).

For LNPs with Cy5-GFP mRNA, DiR dissolved in DMSO was added to the ethanol phase prior to formulation at a final concentration of 0.5 mol%.

siRNA or mRNA was dissolved in 10 mL citrate buffer at pH 3. Each LNP was formulated utilizing an IL-to-RNA weight ratio 10:1 with a volume ratio of aqueous phase to organic phase of 3:1. For the microfluidic-formulated LNPs, each phase was loaded into separate glass syringes (Hamilton Company, Reno, NV, USA) and attached to a Pump 33 DDS syringe pump (Harvard Apparatus, MA, USA). The liquid phases were pushed through a staggered herringbone micromixer microfluidic device, fabricated in polydimethylsiloxane using soft lithography^51^. A two-step exposure process was used to create the SU-8 master with positive channel features on a silicon wafer, where each mixing channel is 4 cm in length. The glass syringes were injected at a flow rate of 0.6 mL/min and 1.8 mL/min for the organic phase and aqueous phase, respectively. Empty LNPs were prepared via the microfluidic devices, but without RNA in the citrate phase. For bulk mixed LNPs, the ethanol and aqueous phases were transferred to a 1.5 mL microcentrifuge tube, and using a repeater pipette, were mixed for 50 cycles at 0.1 mL. All LNPs were dialyzed against pH 7.4 1X PBS in 20 kDa MWCO cassettes for a minimum of 2 h at room temperature. Afterwards, the microfluidic-formulated LNPs were passed through a 220 nm filter and the bulk mixed LNPs were passed through a 450 nm filter, and then were stored at 4 °C.

### Lipid nanoparticle encapsulation and p*K*_a_

Relative encapsulation efficiency and encapsulated RNA concentration were determined via a Quant-iT RiboGreen Assay (Thermo Fisher Scientific). LNPs were diluted 100-fold in 1X tris-EDTA (TE) buffer (Thermo Fisher Scientific) and 1X TE buffer supplemented with 1% (v/v) Triton X-100 (Alfa Aesar, Haverhill, MA, USA). The LNPs equilibrated in the buffers for 5 min before being transferred to a black 96-well plate (Corning, Corning, New York, USA). A standard curve was prepared according to the manufacturer’s instructions using the encapsulated RNA as the standard. The RiboGreen reagent was mixed 1:1 (v/v) with the standard curve and LNP solution and equilibrated for 5 min. Afterwards, fluorescence intensity was read on an Infinite 200 Pro plate reader (Tecan, Morrisville, NC) at an excitation wavelength of 490 nm and an emission wavelength of 530 nm. RNA content was quantified via a standard curve estimated from a univariate least squares linear regression (LSLR). Encapsulation efficiency was calculated as 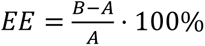, where A is the measured RNA content in TE buffer and B is the measured RNA content post-lysis. The encapsulated RNA concentration was calculated by [𝑐𝑜𝑛𝑐] = 𝐵 − 𝐴.

The relative p*K*_a_ of the LNPs was determined by a 6-(p-toluidinyl)napthalene-2-sulfonic (TNS) assay. LNPs were diluted 3-fold in pH 7.4 1X PBS. Buffers containing 150 mM sodium chloride (MilliporeSigma; St. Louis, MO, USA), 20 mM sodium phosphate (MilliporeSigma), 20 mM ammonium acetate (MilliporeSigma), and 25 mM ammonium citrate (MilliporeSigma) were adjusted from pH 2 to pH 11 in increments of 0.5 pH units. TNS solution was prepared by dissolving TNS (MilliporeSigma) to a concentration of 160 µM in DI water. In a black 96-well plate was added 125 µL of each pH-adjusted solution, 2.5 µL of each LNP, and 5 µL of TNS solution. The plate shook in the dark at 200 rpm for 5 min. Afterwards, fluorescence was measured on a plate reader at an excitation wavelength of 322 nm and an emission wavelength of 431 nm. Using a sigmoidal Four-Parameter Logistic equation, the p*K*_a_ was determined as the pH corresponding to the half-maximum fluorescence intensity (IC50), which corresponds to 50% protonation. Standard deviation was calculated as 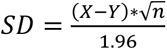, where X is the upper 95% confidence interval, Y is the lower 95% confidence interval, and n is the number of points.

### Dynamic light scattering

Hydrodynamic radius, polydispersity (PDI), and particle concentration were obtained using a DynaPro Plate Reader III (Wyatt Technology, LLC) using a cumulants model. The LNPs were diluted 50-fold in pH 7.4 1X PBS, and 30 µL were loaded onto a 384-well Aurora plate (Wyatt Technology, LLC.). The plate was centrifuged for 5 min at 300 g and then loaded onto the plate reader. Size is reported as intensity-weighted average with *n* = 3 measurements, and data is expressed as mean ± standard deviation, where the standard deviation is calculated by 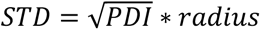.

### Zeta potential

Batch DLS, static light scattering (SLS) and electrophoretic light scattering (ELS) measurements were conducted on a DynaPro ZetaStar^TM^ instrument (Wyatt Technology, LLC.). The instrument is equipped with a 785 nm laser and individually optimized detectors for DLS, SLS, and ELS measurements. The neat LNPs were diluted 100-fold in 0.02 µm filtered 20 mM Tris, pH 7.4, loading 65 µL sample into the ZetaStar dip cell. Data acquisition and analysis were performed with DYNAMICS^TM^ software 8.3.1. (Wyatt Technology, LLC.). Adaptive collection mode was used to optimize the applied current and measurement time. The resulting measurement time was 60 seconds on average.

### Cryogenic transmission electron microscopy

Morphology and size were analyzed by cryo-TEM by adding 3 μL of the LNPs at an mRNA concentration of 50 ng/μL to a Quantifoil^TM^ (Jena, Germany) holey carbon grid that had been glow discharged. Grids were blotted and frozen in liquid ethane using a Vitrobot^TM^ Mark IV (Thermo Fisher Scientific). Imaging was performed at the Beckman Center for cryo-EM on a Titan Krios^TM^ equipped with a K3 Bioquantum (Thermo Fisher Scientific).

### Sedimentation velocity analytical ultracentrifugation

Sedimentation velocity analytical ultracentrifugation (SV-AUC) experiments were performed at 20 °C with an Optima^TM^ analytical ultracentrifuge (Beckman-Coulter, Brea, CA, USA) and a TiAn50 rotor with two-channel charcoal-filled Epon centerpieces and sapphire windows, using both absorbance and interference optics. Data were collected in pH 7.4 1X PBS with detection at 260 and 280 nm, as well as interference optics. Complete sedimentation velocity profiles were recorded every 30 seconds at 20,000 rpm and 20 ^°^C. Data were fit using a least-squares g*(s) model as implemented in the program SEDFIT^52^.

### Asymmetric-flow field-flow fractionation measurements

Samples were injected undiluted in triplicate on an Eclipse^TM^ field flow fractionation (FFF) instrument using a 350 µm fixed-height short channel with dilution control module and a 10 kDa regenerated cellulose membrane (Wyatt Technology, LLC.), connected to a 1260 Infinity II HPLC system with a G1310B isocratic pump and a G1329B autosampler (Agilent Technologies, Inc.). A DAWN^TM^ MALS instrument with an integrated WyattQELS^TM^ DLS detector (Wyatt Technology, LLC.), an Optilab^TM^ differential refractometer (Wyatt Technology, LLC.), and a G1365C UV detector (Agilent Technologies, Inc.) set to 260 nm wavelength were used for online detection. 1X PBS at pH 7.4 was used as the mobile phase. Samples were injected at volumes of 55–150 μL. The FFF system was controlled by VISION 3.2.0 software (Wyatt Technology, LLC.). FFF method parameters consisted of 2.5 mL/min channel flow, 0.5 mL/min detector flow (corresponding to a 5-fold concentration enhancement from the dilution control module), and 0.1 mL/min inject flow. Focusing was at 1.0 mL/ min with a 25% focus position for 14 min. Cross flow was ramped from 0 to 1.0 mL/min in the first minute of the method, and then kept constant at 1.0 mL/min throughout focusing and the initial 5 min of the elution step, followed by an exponential gradient of 1.0–0.04 mL/min over 30 min after which the crossflow was kept constant at 0.04 mL/min for 15 min. Prior to LNP injections, the membrane was conditioned by injecting 100 μL of a 2 mg/mL bovine serum albumin (BSA) standard (Wyatt Technology, LLC.). The system performance was checked by injecting and analyzing a triplicate of 25 μL of the same BSA standard using the BSA short channel FFF method built into the VISION 3.2.0 software. The online MALS detectors were calibrated at a 90° scattering angle and the remaining detector angles normalized to the response at 90° using the BSA monomer peak data. Data acquisition and analysis were performed using the LNP Analysis Module in the ASTRA^TM^ 8.2.2 software (Wyatt Technology, LLC.). Blank injections of mobile phase were used for RI signal baseline subtraction. To obtain accurate concentration data from the online 260 nm UV absorbance signal, empty LNP samples were measured with the same FFF method as the mRNA-LNP samples, and the results from the empty samples were used to generate experimentally derived UV scattering correction profiles with the ASTRA LNP Analysis Module. The scattering correction profiles were then applied non-destructively to the data collected for the corresponding mRNA-LNP samples.

### Synchrotron size-exclusion chromatography in line with small angle X-ray scattering

Size-exclusion chromatography (SEC)-SAXS data were collected at beamline 16-ID (LiX) of the National Synchrotron Light Source II (Upton, NY)^53–55^. SAXS/WAXS data were simultaneously collected at a wavelength of 0.89 Å, yielded accessible scattering angle where 0.005 < q < 3.0 Å^-1^, where q is the momentum transfer, defined as 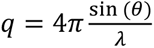, where 𝜆 is the X-ray wavelength and 2𝜃 is the scattering angle; data to q < 0.5 Å^-1^ were used in subsequent analyses. LNPs in volumes of 100 μL and concentrations of ∼10^11^ particles/mL were injected and eluted isocratically at 0.35 ml/min from a prepared 3 mL Sepharose 4B (MilliporeSigma) column or TOSOH TSKgel G6000PWXL-CP SEC 7.8 mm × 300 mm and 13 µm particle size column (MilliporeSigma) equilibrated in pH 7.4 1X PBS at room temperature. Eluent from the column flowed into a 1 mm capillary for subsequent X-ray exposures at 2-s intervals. Plots of intensity from the forward scatter closely correlated to in-line UV and refractive index (RI) measurements.

### Small angle X-ray scattering analysis

Analysis of the SEC-SAXS data sets were performed in the program RAW^56^. Buffer subtracted profiles were analyzed by singular value decomposition (SVD) and the ranges of overlapping peak data were determined using evolving factor analysis (EFA) as implemented in REGALS^57^. The determined peak windows were used to identify the basis vectors for each component and the corresponding SAXS profiles were calculated. When manually fitting the pair distribution function P(r), the maximum diameter of the particle (D_max_) was incrementally adjusted in GNOM^58^ as implemented in RAW^59^ to maximize the Total Estimate and χ^2^ figures, to minimize the discrepancy between the fit and the experimental data to a q_max_ of 0.1 Å^-1^, and to optimize the visual qualities of the distribution profile.

Deconvolution of the primary Braggs peaks in SAXS were using multiple Lorentz fits with the built-in function:

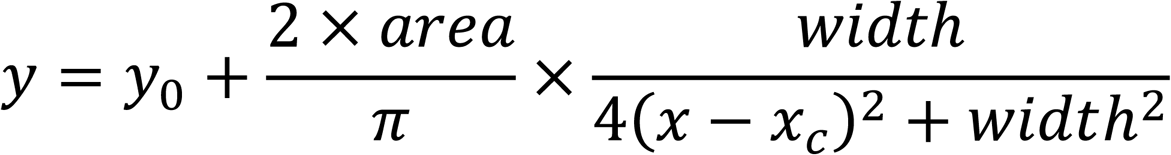

where y_0_ is an offset and was set to the X-ray baseline and x_c_ is a center of function and corresponds to the center of the SAXS peak.

Porod values were calculated using ScÅtter. The Porod region of the primary scattering profiles is defined by the region before the peak structural features (q < 0.1 Å^-1^) and after the Guinier region (q > 0.01 Å^-1^); the scattering in this region decays as I(q) ∝ q^-P^, where P is the Porod exponent that is dependent on particle shape^60^.

DENSS^61^ was used to calculate the *ab initio* electron density map directly from the GNOM output. Twenty reconstructions of electron density were performed in the slow mode with default parameters and subsequently averaged and refined. Reconstructions were visualized using PyMOL 2.5.2 Molecular Graphics System (Schrodinger, LLC, New Your, NY) with five contour levels of density rendered with these respective colors: 15σ (red), 10σ (green), 5σ (cyan), 2.5σ (blue), and -0.7σ (blue). The sigma (σ) level denotes the standard deviation above the average electron density value of the generated model.

### *In vitro* transfection

Human primary T cells were obtained from donors at the Human Immunology Core at the University of Pennsylvania. T cells were activated using Human T-Activator CD3/CD28 Dynabeads^TM^ (Thermo Fisher Scientific) at a bead-to-cell ratio of 1:1 for 24 h. T cells were cultured in Roswell Park Memorial Institute (RPMI; Gibco, Dublin, Ireland) medium supplemented with 10% fetal bovine serum (FBS; Gibco) and 1% penicillin-streptomycin (P/S; Gibco). T cells were plated onto white 96-well plates (Corning) at a density of 60,000 cells per well. Immediately afterwards, the cells were treated with LNPs containing FLuc mRNA at a dose of 300 ng per well. Following 24 h of incubation, the plates were centrifuged at 400 g for 5 min to pellet the cells. Then, luminescence and viability were ascertained via a Promega (Madison, WI, USA) Luciferase Assay System and a Cell-titer Glo® Luminescent Cell Viability Assay, respectively, according to the manufacturer’s instructions.

FaDu cells were obtained from ATCC (HTB-43) and cultured in Dulbecco’s modified Eagle medium (DMEM; Gibco) with L-glutamine supplemented with 10% FBS and 1% P/S. The FaDu cells were plated onto white 96-well plates at a density of 20,000 cells per well and allowed to adhere for 16 h. Afterwards, the cells were treated with LNPs containing FLuc mRNA at a dose of 20 ng per well and luminescence and viability were quantified as described above.

### Murine biodistribution

C57BL/6J female mice of 6–8 weeks old with an average weight of 20 g were purchased from Jackson Laboratory (Bar Harbor, ME). Animals were housed in a barrier facility with air humidity 40%–70%, ambient temperature (22 ± 2 °C), and 12 h dark/12 h light cycle.

For intravenous studies, mice were injected with LNPs encapsulating FLuc mRNA via the lateral tail vein at a dose of 0.1 mg of mRNA per kg of body mass (mg/kg). For intramuscular studies, mice were injected in the left quadricep at a dose of 0.05 mg/kg. After 6 h, the mice were administered an intraperitoneal injection of D-luciferin (0.2 mL, 15 mg/mL; Biotium, Fremont, CA). Then, after 5 min, full body luminescence images were obtained using an In Vivo Imaging System (IVIS; PerkinElmer, Waltham, MA) for the intramuscular group only. For the intravenous group, the mice were euthanized, and the heart, lungs, liver, kidneys, and spleen were removed and imaged for luminescence using IVIS. Total flux was quantified by the Living Image^TM^ Software 4.7.3 (PerkinElmer) by placing rectangular region of interests (ROI) around the full body or organ images, keeping the same ROI sizes among each body or organ.

### Erythropoietin toxicity

C57BL/6J female mice of 6–8 weeks old with an average weight of 20 g were intravenously administered 5moU LNPs encapsulating epo mRNA at 0.4 mpk. Mice were monitored for 1 h. After 6 h from the injection, blood was collected via retro-orbital bleeding in Microtainer blood collection tubes containing serum separator gel (BD, Franklin Lakes, NJ, USA). Blood was centrifuged for 15 min at 2,000 g and serum was isolated. Epo levels were obtained using a Human Erythropoietin ELISA (Thermo Fisher Scientific). TNF-α and IL-6 levels were measured using BioLegend (San Diego, CA, USA) ELISA Max^TM^ Deluxe Set Murine TNF-α and IL-6 using Nunc^TM^ MaxiSorp^TM^ ELISA plates, uncoated (BioLegend). Aspartate transaminase (AST) values were obtained using a colorimetry activity assay (Cayman Chemical Company, Ann Arbor, MI, USA). Alanine transaminase (ALT) levels were obtained using a mouse ALT ELISA (Abcam, Cambridge, UK). All ELISAs and colorimetric assays were performed according to the manufacturer’s instructions.

### Endocytosis studies

Human primary T cells (1:1, CD4+ and CD8+) activated as described above, were treated with DiR-dyed LNPs encapsulating 5moU Cy5-GFP. T cells were treated with 600 ng of LNP per 250,000 cells in 24-well clear bottom plates (Corning). After 24 h, the Dynabeads were removed via a magnet, and the cells were centrifuged for 5 min at 400 g. The media was removed, and the cells were resuspended in pH 7.4 1X PBS. Flow cytometry was performed on a BD FACSymphony A5 SE Flow Cytometer (Becton Dickinson, Franklin Lakes, NJ, USA) and analyzed via FlowJo V10.10 (Becton Dickinson).

For the endocytosis inhibitor studies, T cells were incubated with 2 mM amiloride (Thermo Fisher Scientific), 500 µM methyl-β-cyclodextrin (Thermo Fisher Scientific), 2 µM chlorpromazine (MilliporeSigma), 100 µM Dynasore (MilliporeSigma), or 200 µM Genistein (MilliporeSigma) for 30 min. The cells were then centrifuged for 5 min at 400 g, where the supernatant was aspirated and the cells were resuspended in fresh supplemented RPMI. The cells were then treated with FLuc LNPs at a dose of 300 ng per 60,000 cells. After 24 h, luciferase and toxicity assays were conducted as described above. For luminescence, each group was normalized to T cells, which were not incubated with endocytosis inhibitor but treated with the corresponding LNP. For toxicity, no LNPs were added, and each group was normalized to T cells that were not incubated with endocytosis inhibitors.

### Statistics & reproducibility

All statistical analysis was performed in GraphPad^TM^ Prism Version 10.4.0 (GraphPad Software, Inc, La Jolla, CA, USA). All tests of significance were performed at a significance level of α = 0.05. For experiments with one variable and multiple technical or biological replicates, one-way analyses of variance (ANOVAs) with post hoc Holm-Šídák correction for multiple comparisons were used to compare responses across treatment groups. For experiments that measured two variables with more than two treatment groups, two-way ANOVAs with post hoc Holm-Šídák correction for multiple comparisons were used. For experiments that measured two variables with two treatment groups in each variable, two-sided multiple unpaired t tests with Holm-Šídák correction for multiple comparisons were used. All data are presented as mean ± standard deviation unless otherwise reported.

## Data availability

All data supporting the findings of this study are available within the paper and Supplementary Information. Source data for traditional characterization methods and biological testing are provided as a Source File. The SV-AUC, FFF-MALS, and SEC-SAXS raw and analyzed data are available at https://10.5281/zenodo.17042311.

## Code availability

No original code was generated for this study.

