## Supplemental Information for "Elucidating lipid nanoparticle properties and structure through biophysical analyses"

### Supplementary Information

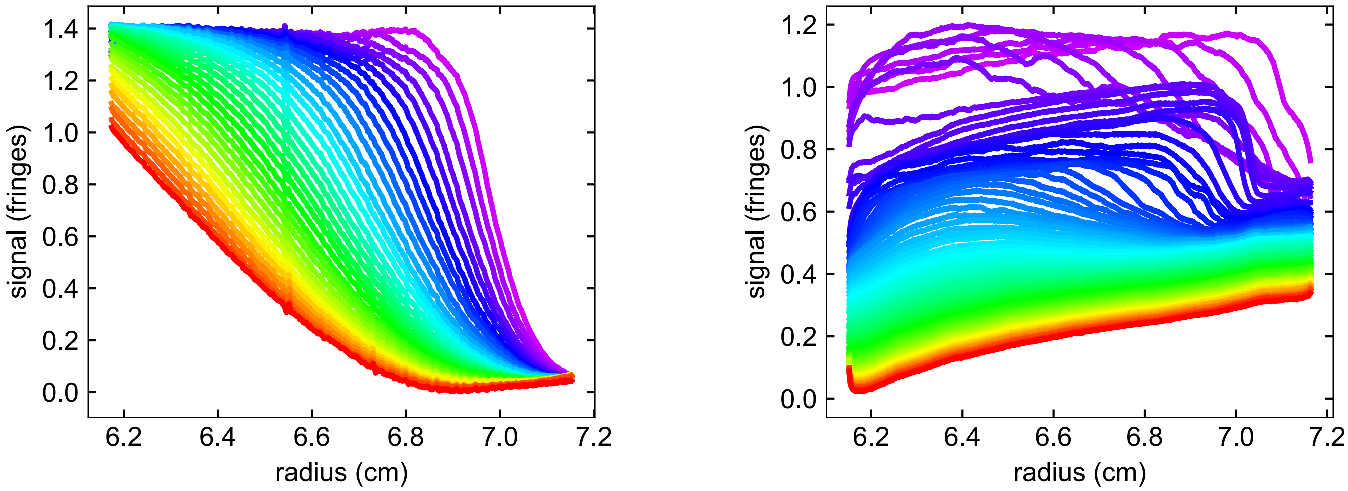


**Supplemental Figure 1. Representative sedimentation velocity analytical ultracentrifugation (SV-AUC) data.** Shown are the experimental data for MC3$^{\mu F}$ with no mRNA (left) and MC3$^{\mu F}$ encapsulating firefly luciferase mRNA (right). Both data show evidence for flotation (boundary migration from right to left), and the mRNA-containing sample additionally shows evidence of sedimentation. These figures were generated using the program GUSSI[1]. Source data are provided in Zenodo (10.5281/zenodo.17042311).

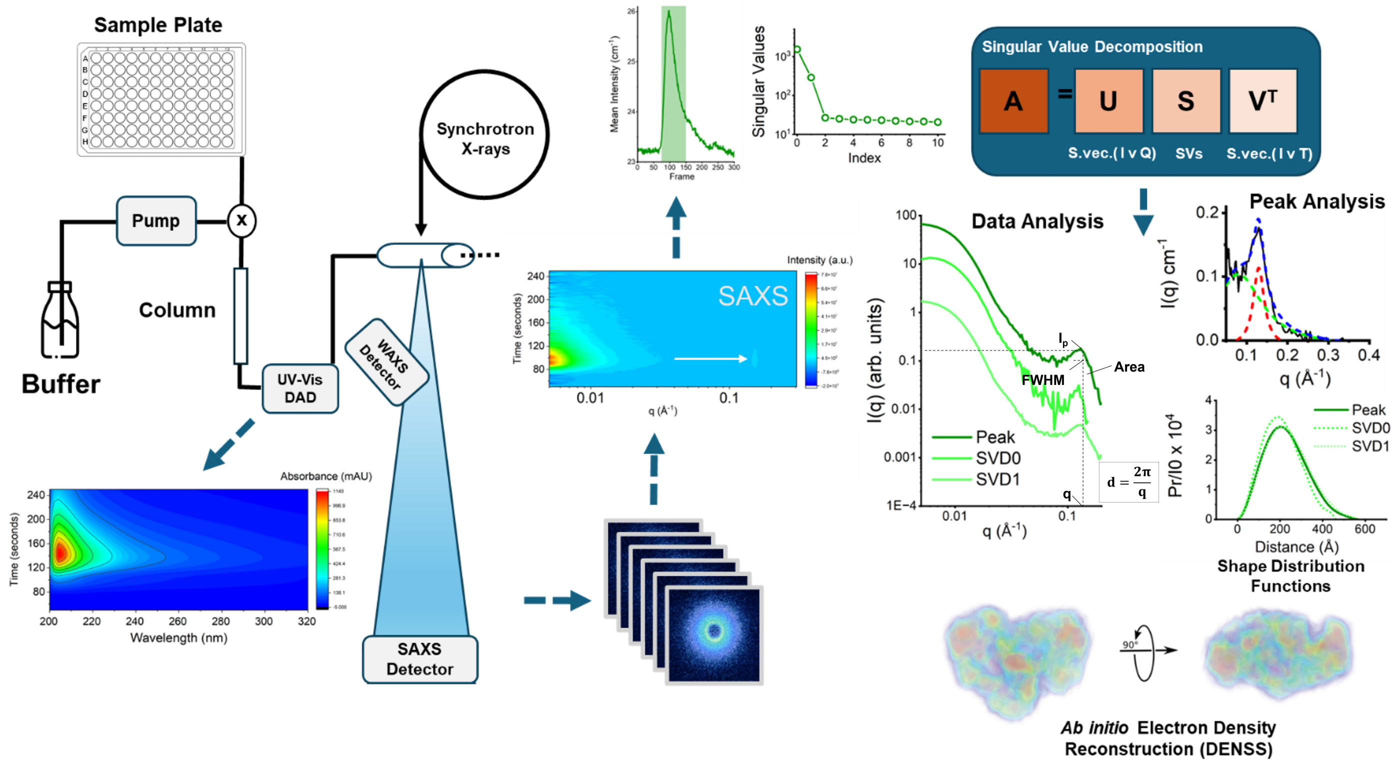


**Supplementary Figure 2. Overview SEC-SAXS workflow.** LNPs isocratically eluted from a size-exclusion column are analyzed via a UV/Vis diode array detector preceding collection of synchrotron small-angle X-ray scattering on the eluant. The chromatogram from the mean X-ray intensity represents the collection of scattering profiles (~300 profiles). A region of this dataset is selected for single value decomposition (SVD) to extract individual species that can then be further processed to ascertain the peak features and shape.

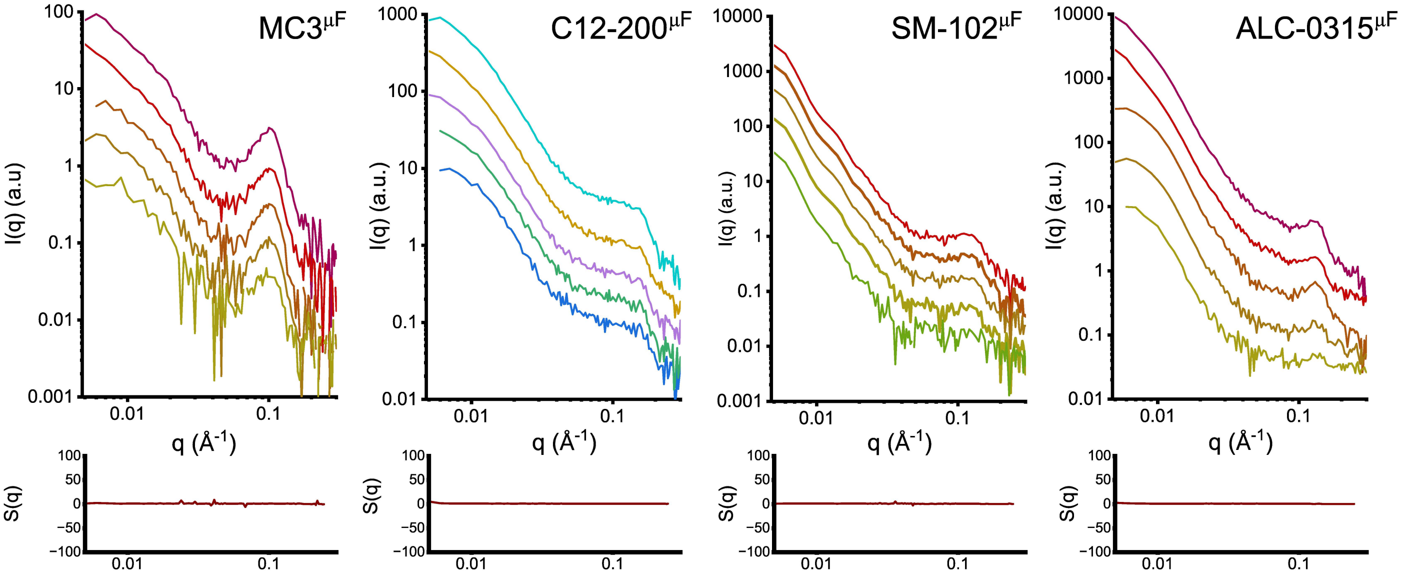


**Supplemental Figure 3. Concentration series analysis using batch SAXS measurements reveals no evidence of interparticle interactions from S(q).** Shown in each upper panel are concentration series for each microfluidic mRNA-LNP, in a log-log plot where $0.005 < q < 0.3$ Å$^{-1}$. In the lower panel for each particle a plot of S(q) is shown, from the ratio of the highest and lowest concentration, with the assumption that the lowest concentration examined properly represents the form factor FF(q). From this analysis, no evidence of an interaction parameter is apparent. Source data are provided in Zenodo (10.5281/zenodo.17042311).

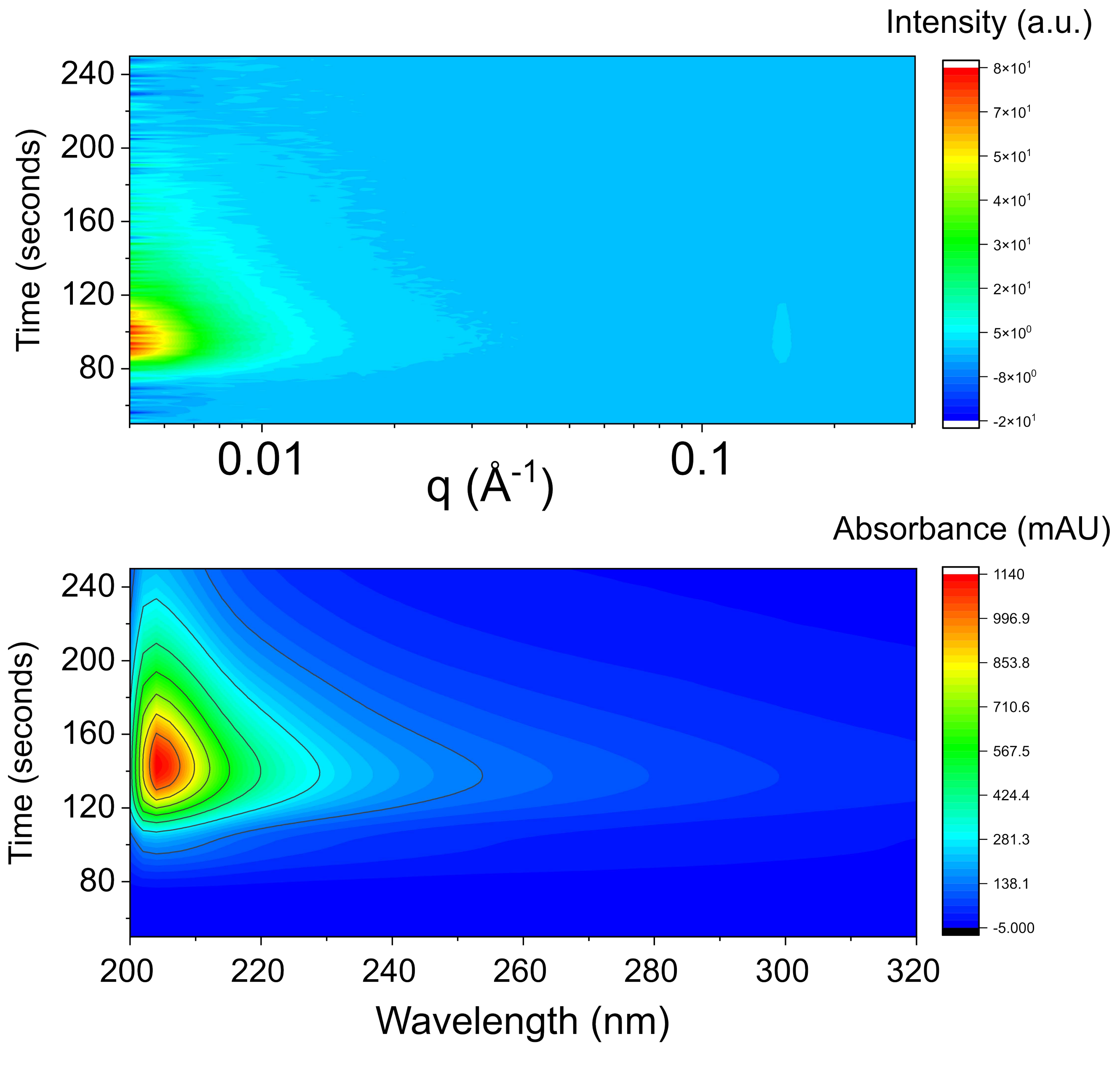


**Supplemental Figure 4. Representative X-ray and UV diode array detector data from SEC-SAXS analysis.** Shown in the upper panel as a heat map is the intensity of scattering ($I_q$) as a function of scattering angle q and time where $0.005 < q < 0.3$ Å$^{-1}$; only the first four minutes of the ten-minute method are shown. Denoted with an arrow is the intensity corresponding to the first-order Bragg peak which was used to identify RNA containing species for subsequent SVD analysis. In the lower panel shown is UV absorbance data as a heat map, as a function of both wavelength and time. The strongest 260 nm signal correlating to the presence of RNA correlates with SAXS when accounting for inter-detector delay. Source data are provided in Zenodo (10.5281/zenodo.17042311).

**Supplementary Table 1: Static SAXS parameters.**

| LNP | Method | q (Å$^{-1}$)$^{Static}$ | $I_p$ (cm$^{-1}$)$^{Static}$ | d (Å)$^{Static}$ | Area$^{Static}$ | FWHM$^{Static}$ | $R_g$ (Å)$^{Static}$ | $D_{max}$ (Å)$^{Static}$ |
|---|---|---|---|---|---|---|---|---|
| MC3 | µF | 0.10 | 0.13 | 62.8 | 0.00780 | 0.0372 | 173 | 488 |
| | bulk | *n.d.* | *n.d.* | *n.d.* | *n.d.* | *n.d.* | *n.d.* | *n.d.* |
| C12-200 | µF | 0.16 | 0.03 | 40.5 | 0.00190 | 0.0408 | 181 | 562 |
| | bulk | *n.d.* | *n.d.* | *n.d.* | *n.d.* | *n.d.* | *n.d.* | *n.d.* |
| SM-102 | µF | 0.13 | 0.01 | 48.3 | 0.00004 | 0.0033 | *n.a.* | *n.a.* |
| | bulk | *n.d.* | *n.d.* | *n.d.* | *n.d.* | *n.d.* | *n.d.* | *n.d.* |
| ALC-0315 | µF | 0.13 | 0.04 | 48.3 | 0.00170 | 0.0296 | *n.a.* | *n.a.* |
| | bulk | *n.d.* | *n.d.* | *n.d.* | *n.d.* | *n.d.* | *n.d.* | *n.d.* |

**Supplementary Table 2: Average peak parameters from SEC-SAXS.**

| LNP | Method | q (Å$^{-1}$)$^{Avg}$ | $I_p$ (cm$^{-1}$)$^{Avg}$ | d (Å)$^{Avg}$ | Area$^{Avg}$ | FWHM$^{Avg}$ | $R_g$ (Å)$^{Avg}$ | $D_{max}$ (Å)$^{Avg}$ | SVD | Porod Values |
|---|---|---|---|---|---|---|---|---|---|---|
| MC3 | µF | 0.097 | 0.09 | 64.7 | 0.0029 | 0.032 | 167 | 475 | 3 | 3.3 |
| | bulk | 0.110 | 0.04 | 57.2 | 0.0011 | 0.025 | *n.a.* | *n.a.* | 2 | 2.4 |
| C12-200 | µF | 0.151 | 0.10 | 41.6 | 0.0021 | 0.019 | 177 | 570 | 3 | 3.1 |
| | bulk | 0.150 | 0.34 | 41.9 | 0.0080 | 0.014 | *n.a.* | *n.a.* | 2 | 2.9 |
| SM-102 | µF | 0.121 | 0.12 | 51.9 | 0.0075 | 0.031 | 169 | 495 | 2 | 3.5 |
| | bulk | 0.125 | 0.07 | 50.4 | 0.0016 | 0.015 | *n.a.* | *n.a.* | 3 | 3.0 |
| ALC-0315 | µF | 0.130 | 0.11 | 48.3 | 0.0063 | 0.033 | 175 | 566 | 2 | 3.6 |
| | bulk | 0.135 | 0.03 | 46.5 | 3.75E-04 | 0.008 | *n.a.* | *n.a.* | 2 | 3.7 |

**Supplementary Table 3: SVD0 parameters from SEC-SAXS.**

| LNP | Method | q (Å$^{-1}$)$^{SVD0}$ | $I_p$ (cm$^{-1}$)$^{SVD0}$ | d (Å)$^{SVD0}$ | Area$^{SVD0}$ | FWHM$^{SVD0}$ | $R_g$ (Å)$^{SV0}$ | $D_{max}$ (Å)$^{SV0}$ | Porod Values |
|---|---|---|---|---|---|---|---|---|---|
| MC3 | µF | 0.094 | 0.033 | 66.8 | 0.0011 | 0.026 | 197 | 556 | *n.a.* |
| | bulk | 0.104 | 0.009 | 60.4 | 2.45E-04 | 0.021 | *n.a.* | *n.a.* | 3.1 |
| C12-200 | µF | 0.150 | 0.113 | 41.9 | 0.0018 | 0.020 | 191 | 597 | 2.9 |
| | bulk | 0.150 | 0.111 | 41.9 | 0.0023 | 0.013 | *n.a.* | *n.a.* | 2.9 |
| SM-102 | µF | 0.120 | 0.019 | 52.3 | 0.0017 | 0.038 | 179 | 499 | *n.a.* |
| | bulk | 0.125 | 0.002 | 50.2 | 1.68E-04 | 0.013 | *n.a.* | *n.a.* | *3.5* |
| ALC-0315 | µF | 0.125 | 0.003 | 50.2 | 4.30E-04 | 0.021 | 153 | 461 | 3.5 |
| | bulk | 0.135 | 9.99E-04 | 46.5 | 1.67E-04 | 0.245 | *n.a.* | *n.a.* | 3.8 |

**Supplementary Table 4: SVD1 parameters from SEC-SAXS.**

| LNP | Method | q (Å$^{-1}$)$^{SVD1}$ | $I_p$ (cm$^{-1}$)$^{SVD1}$ | d (Å)$^{SVD1}$ | Area$^{SVD1}$ | FWHM$^{SVD1}$ | $R_g$ (Å)$^{SV1}$ | $D_{max}$ (Å)$^{SV1}$ | Porod Values |
|---|---|---|---|---|---|---|---|---|---|
| MC3 | μF | 0.098 | 0.032 | 64.1 | 0.0017 | 0.047 | 145 | 412 | *n.a.* |
| | bulk | n.d. | n.d. | n.d. | n.d. | n.d. | *n.a.* | *n.a.* | 3.1 |
| C12-200 | μF | 0.148 | 0.039 | 42.4 | 0.0019 | 0.039 | 140 | 419 | 2.9 |
| | bulk | n.d. | n.d. | n.d. | n.d. | n.d. | *n.a.* | *n.a.* | 2.9 |
| SM-102 | μF | 0.120 | 0.027 | 52.3 | 0.0012 | 0.030 | 170 | 503 | 3.5 |
| | bulk | 0.124 | 0.01 | 50.6 | 2.01E-04 | 0.014 | *n.a.* | *n.a.* | 2.6 |
| ALC-0315 | μF | 0.129 | 0.018 | 48.7 | 7.90E-04 | 0.034 | 180 | 554 | 3.5 |
| | bulk | 0.132 | 0.015 | 47.5 | 9.18E-04 | 9.36E-05 | *n.a.* | *n.a.* | 3.9 |

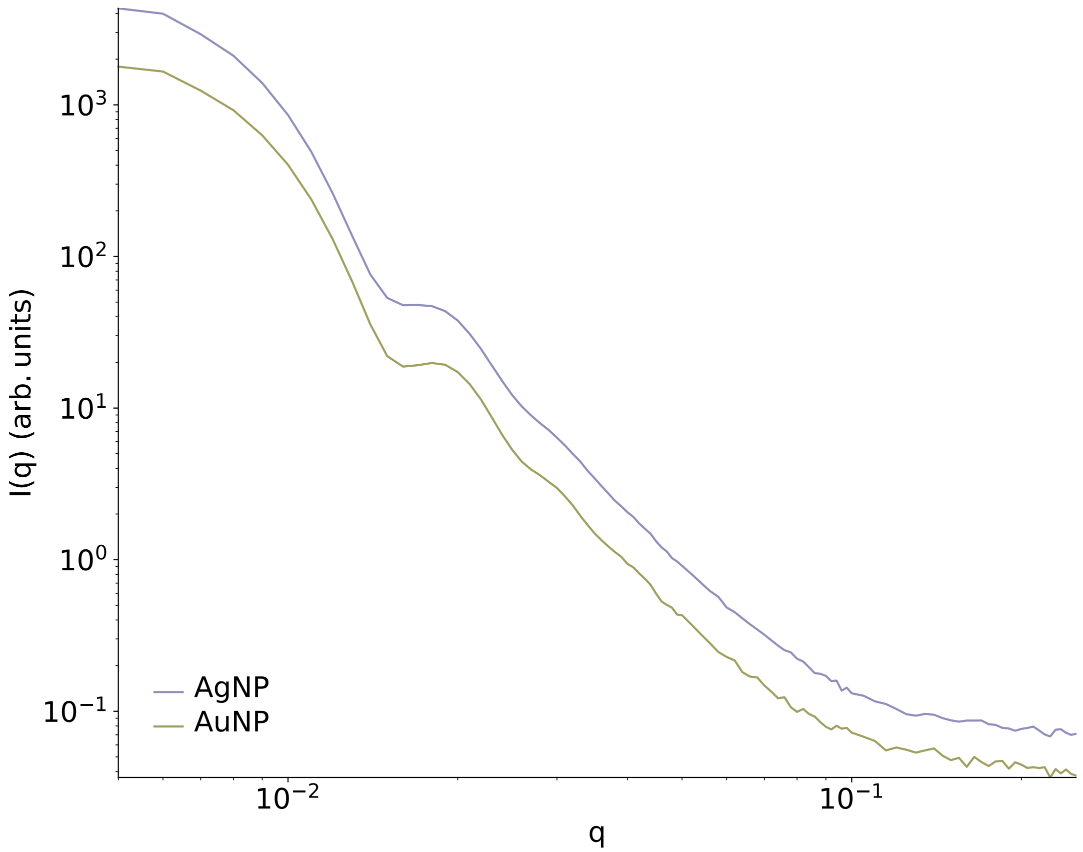


**Supplementary Figure 5: SEC-SAXS analysis of control particles.** SAXS profiles of 60 nm gold and silver spherical nanoparticles. Source data are provided in Zenodo (10.5281/zenodo.17042311).

**Supplementary Table 5: Shape factor ($\frac{D_{max}}{R_g}$) from P(r) analysis.**

| | | |
|---|---|---|
| $MC3^{\mu F}$ | Average | 2.844 |
| | SVD0 | 2.822 |
| | SVD1 | 2.841 |
| $C12\text{-}200^{\mu F}$ | Average | 3.220 |
| | SVD0 | 3.126 |
| | SVD1 | 2.993 |
| $SM\text{-}102^{\mu F}$ | Average | 2.929 |
| | SVD0 | 2.788 |
| | SVD1 | 2.959 |
| $ALC\text{-}0315^{\mu F}$ | Average | 3.234 |
| | SVD0 | 3.013 |
| | SVD1 | 3.078 |

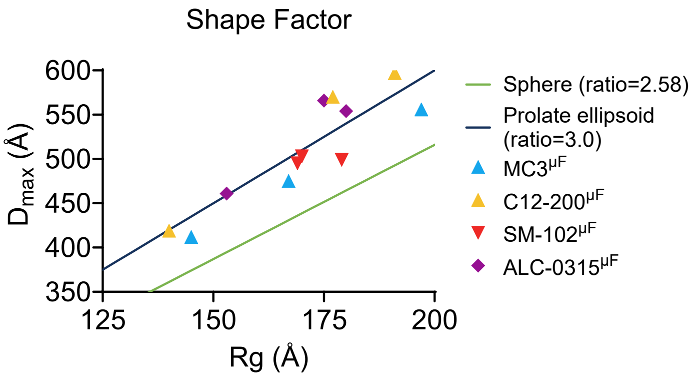


**Supplementary Figure 6: Shape factor analysis**. Plots of $D_{max}$ vs $R_g$ for each microfluidic LNP, obtained through P(r) analysis. For each LNP, the values for the average, SVD0, and SVD1 P(r) analyses are plotted. Solid lines representing sphere and prolate ellipsoid shape factors are plotted as reference for each $D_{max}$ and $R_g$ value.

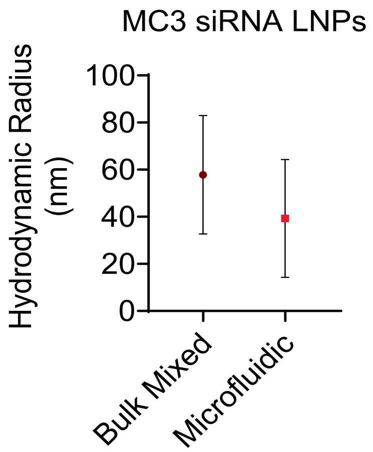


**Supplementary Figure 7: Dynamic light scattering analysis of siRNA LNPs.** Hydrodynamic radii of bulk mixed and microfluidic formulated MC3 LNPs encapsulating siRNA. Data are reported as mean ± SD of *n* = 3. Source data are provided as a Supplementary Source Data file.

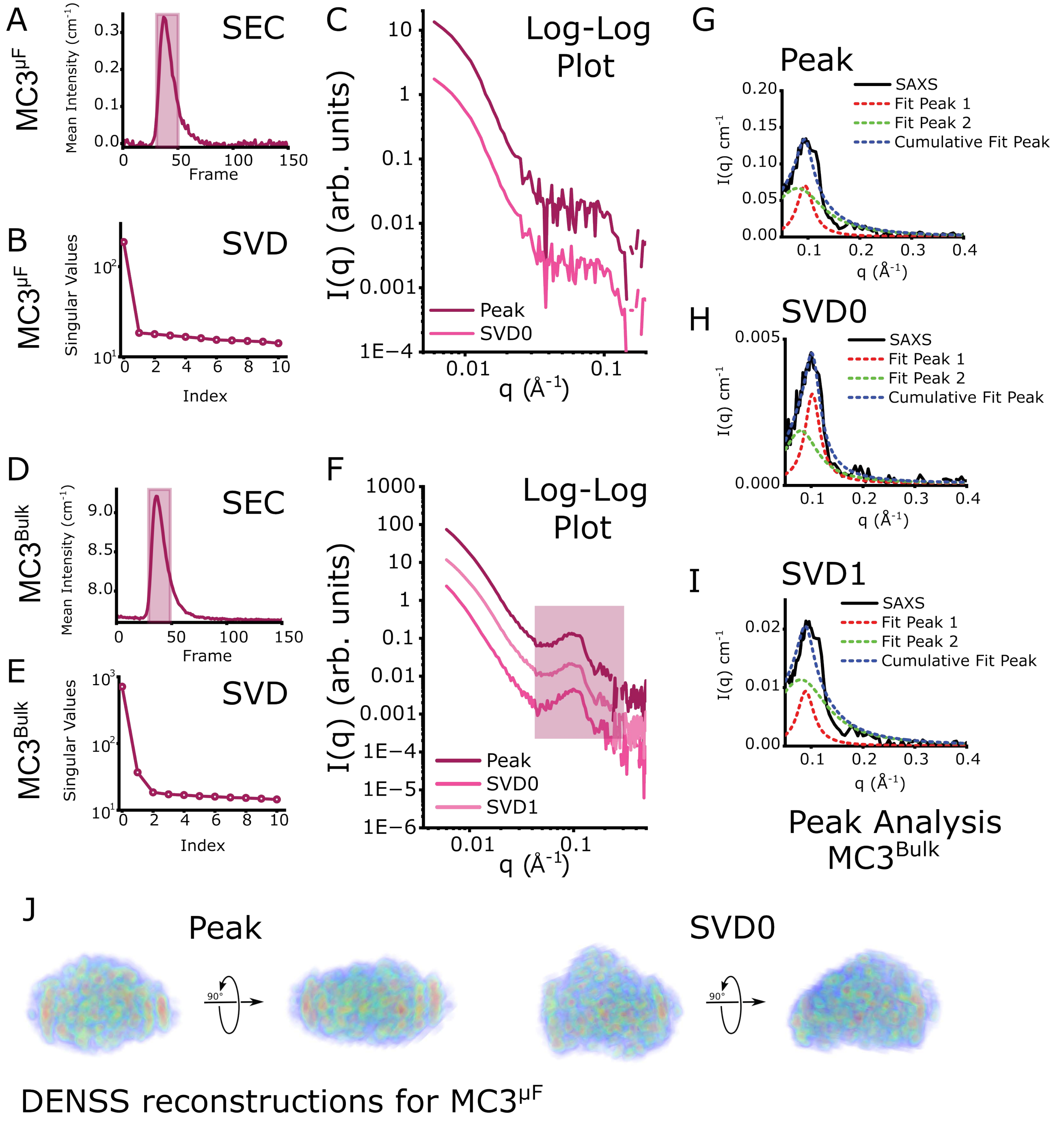


**Supplementary Figure 8: SEC-SAXS analysis of MC3 siRNA LNPs. A–C,** MC3$^{\mu F}$ siRNA LNP analysis via SEC-SAXS, involving **(A)** identification of the main LNP peak in the SEC chromatogram, **(B)** selection of the number of singular values and **(C)** scattering intensity of the main peak and corresponding singular value. **D–F,** MC3$^{Bulk}$ siRNA LNP analysis via SEC-SAXS, involving **(D)** identification of the main LNP peak in the SEC chromatogram, **(E)** selection of the number of singular values and **(F)** scattering intensity of the main peak and corresponding singular values. **G–I,** Fitting of the higher-q peak features for the MC3$^{Bulk}$ siRNA LNP using multiple Lorentz peak feature for **(G)** the peak average, **(H)** SVD0, and **(I)** SVD1, where red is the first order Braggs peak fit, green represents higher-order particle disorder, and blue is the cumulative fit of the two features. **J,** DENSS *ab initio* electron density reconstructions from the SEC-SAXS profiles of the MC3$^{\mu F}$ siRNA LNP for the average peak and SVD0. Source data are provided in Zenodo (10.5281/zenodo.17042311).

**Supplementary Table 6. Two-tailed P values for Spearman correlations.**

| Parameter | T cells | Intravenous | Intramuscular |
|---|---|---|---|
| $I_p(cm^{-1})^{Avg}$ | 0.0368 | 0.0368 | 0.793 |
| $I_p(cm^{-1})^{SVD0}$ | 0.0368 | 0.9349 | 0.1150 |
| $I_p(cm^{-1})^{SVD1}$ | 0.1750 | 0.6583 | 0.1028 |
| $q(Å^{-1})^{Avg}$ | >0.9999 | 0.2675 | 0.7520 |
| $q(Å^{-1})^{SVD0}$ | 0.8401 | 0.4279 | 0.882 |
| $q(Å^{-1})^{SVD1}$ | 0.1750 | 0.9194 | 0.4972 |
| $d(Å)^{Avg}$ | >0.9999 | 0.2675 | 0.7520 |
| $d(Å)^{SVD0}$ | 0.9248 | 0.2560 | 0.8506 |
| $d(Å)^{SVD1}$ | 0.1750 | 0.9194 | 0.4972 |
| $Area^{Avg}$ | 0.0218 | 0.0576 | 0.7033 |
| $Area^{SVD0}$ | 0.0154 | 0.3599 | 0.3894 |
| $Area^{SVD1}$ | 0.2417 | 0.3556 | 0.0583 |
| $FWHM^{Avg}$ | 0.7520 | 0.7930 | >0.9999 |
| $FWHM^{SVD0}$ | 0.3907 | 0.4278 | 0.9248 |
| $FWHM^{SVD1}$ | 0.1361 | 0.7139 | 0.2417 |
| Nucleic acid number | 0.2992 | 0.8401 | 0.3894 |
| Lipid concentration | 0.5821 | 0.7033 | 0.2992 |
| $M_w/M_n$ | 0.0831 | 0.5008 | 0.2431 |
| $M_w$ | 0.9768 | 0.7520 | 0.5008 |
| $R_{geometric}$ | 0.9768 | 0.8401 | 0.5821 |
| $R_{h,MALS}$ | 0.9349 | 0.7520 | 0.8401 |
| $R_{h,DLS}$ | 0.8820 | 0.8401 | 0.5364 |
| $PDI_{DLS}$ | 0.9349 | 0.1150 | 0.5364 |
| Relative p$K_a$ | 0.8820 | 0.3599 | 0.0022 |
| Encapsulation efficiency | 0.115 | 0.6646 | 0.793 |
| ζ-potential | >0.9999 | 0.7033 | 0.6191 |

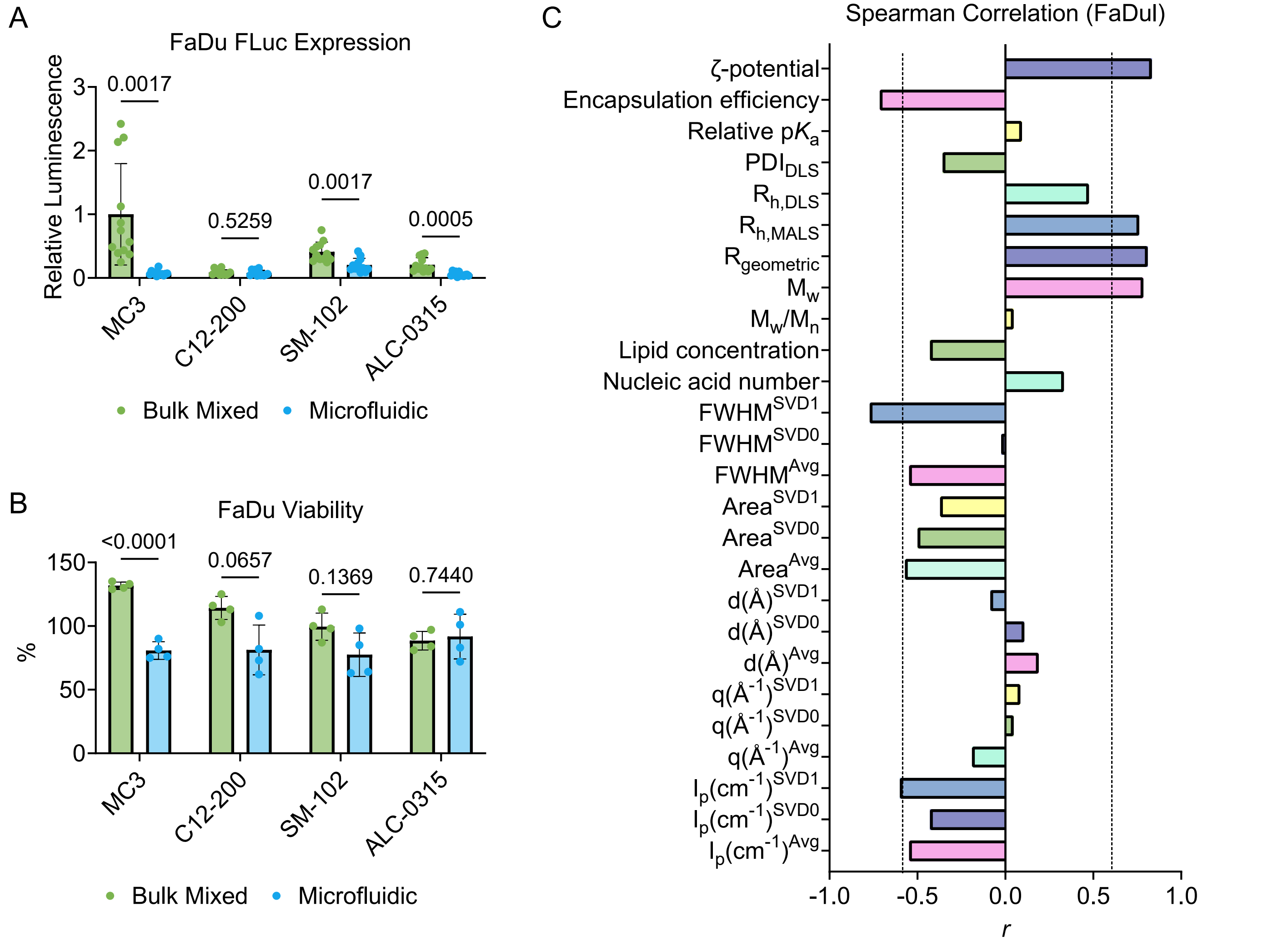


**Supplementary Figure 9: LNP transfection in FaDu cells and correlations. A–B,** FaDu cells were plated at 20,000 cells per well and after adhering for 16 h, were treated with the eight LNPs at a dose of 20 ng of encapsulated FLuc mRNA per well. After 24 h, **(A)** luminescence and **(B)** toxicity were quantified. Relative luminescence and viability are reported as mean ± SD of $n$ = 12 for luminescence and $n$ = 4 for viability. **D,** Spearman correlations for the relative luminescence values utilizing the physicochemical parameters from the traditional, FFF-MALS, and SEC-SAXS methods. **A–B,** Two-sided multiple unpaired T test with *post hoc* Holm–Šídák correction for multiple comparisons was used to compare bulk mixed against microfluidic-formulated LNP **(A)** relative luminescent and **(B)** toxicity values for each LNP group. Source data are provided as a Supplementary Source Data file.

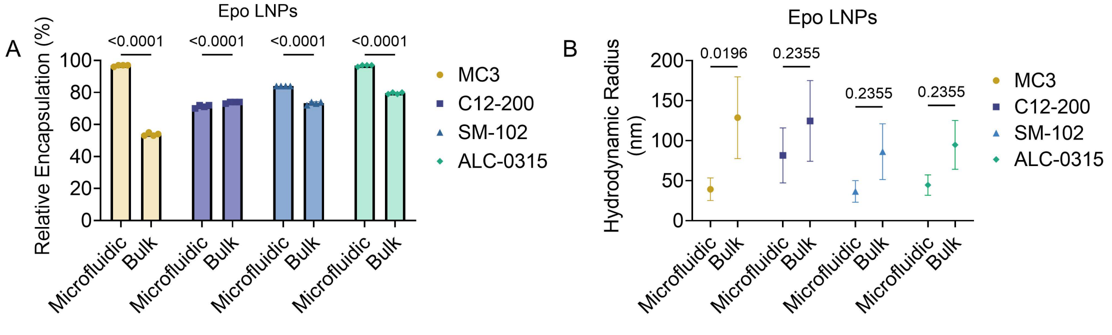


**Supplementary Figure 10: Epo LNP physicochemical characteristics. A–B,** LNP traditional characterization of **(A)** relative mRNA encapsulation efficiency determined by RiboGreen assay and **(B)** hydrodynamic radius obtained via DLS. Measurements are reported as mean ± SD of $n$ = 4 technical replicates for encapsulation efficiency and $n$ = 3 technical replicates for hydrodynamic radius. Two-way ANOVA with Holm–Šídák correction for multiple comparisons were used to compare the **(A)** relative encapsulation efficiency and **(B)** hydrodynamic radius across formulation techniques. Source data are provided as a Supplementary Source Data file.

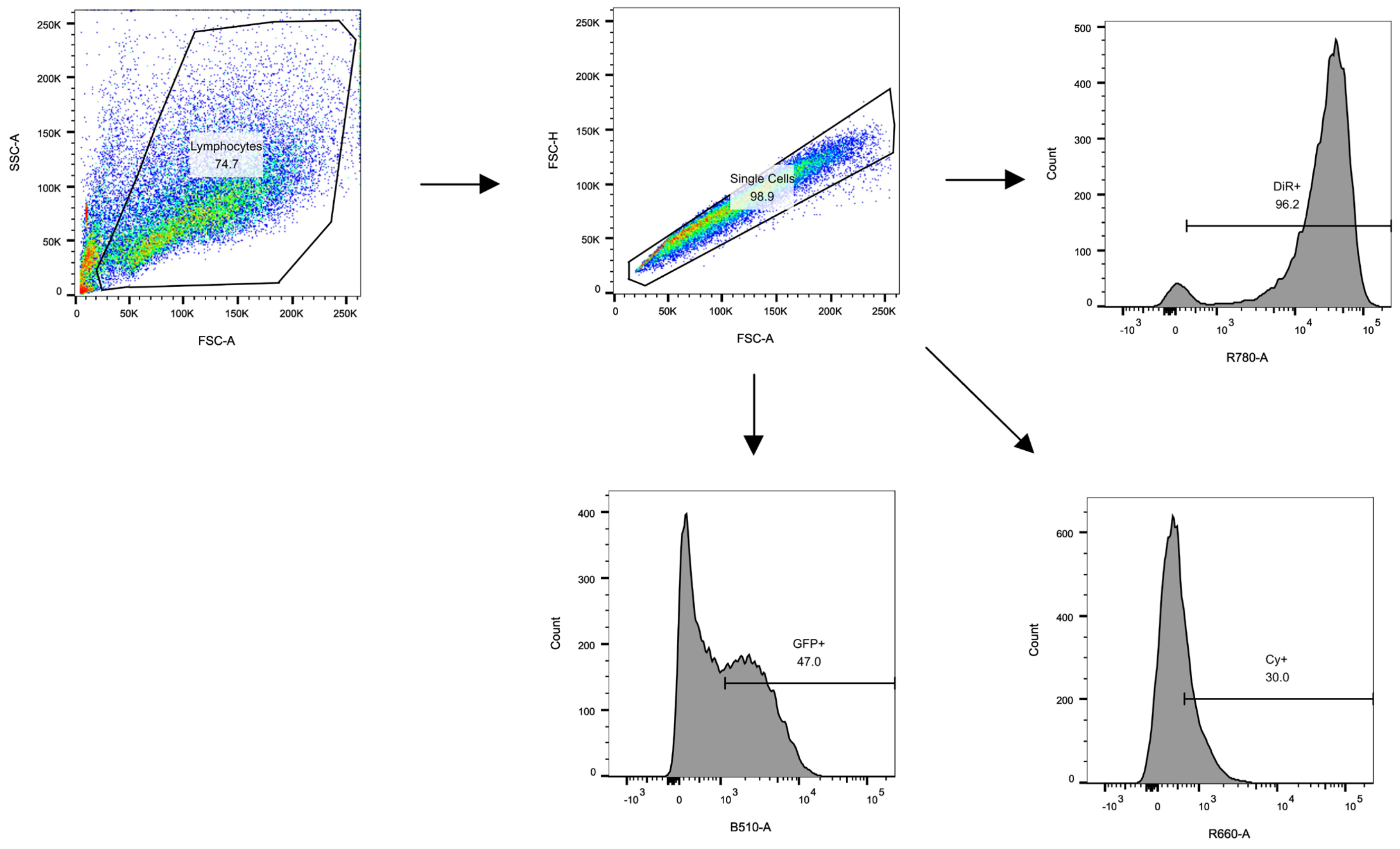


**Supplementary Figure 11: Gating strategy for flow cytometry analysis of T cells.** Gating strategy to examine DiR (R780), Cy5 (R660), and GFP (B510) from primary human T cells treated with LNPs containing 0.5 mol% DiR and Cy5-tagged GFP mRNA.

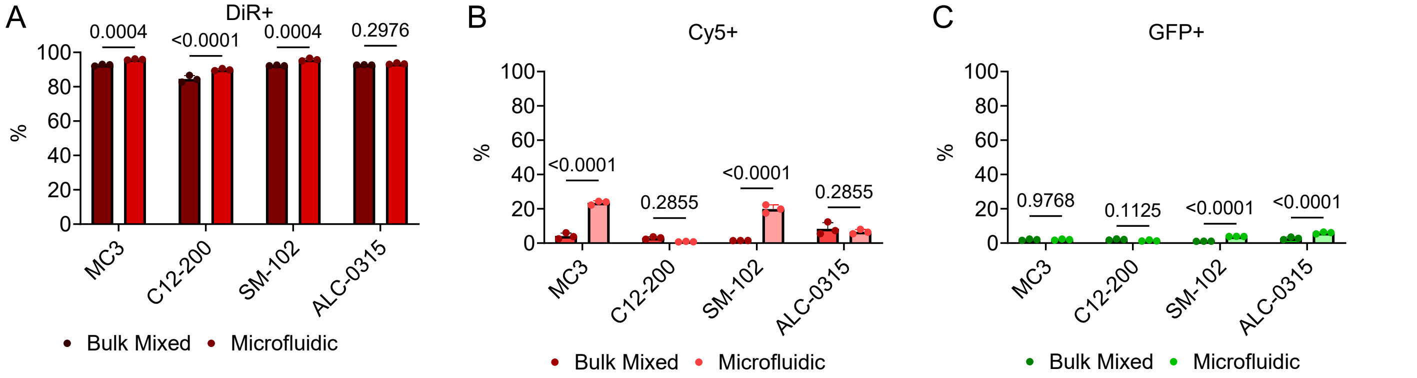


**Supplementary Figure 12: Low dose LNP flow experiment in T cells. A–C,** Human primary T cells (1:1, CD4+/CD8+) were activated for 16 h with human CD3/CD28 Dynabeads$^{TM}$ (1:1, bead/cell). Afterwards, LNPs containing 0.5 mol% DiR and Cy5-mRNA were added to T cells at an mRNA concentration of 50 ng per 250,000 cells. After 24 h, the beads were removed and the cells were analyzed by flow cytometry to ascertain the percentage of cells with **(A)** DiR, **(B)** Cy5, and **(C)** GFP. Positive cell percentage is reported as mean ± SD of $n$ = 3. **A–C,** Two-sided multiple unpaired T test with *post hoc* Holm–Šídák correction for multiple comparisons was used to compare bulk mixed against microfluidic-formulated LNPs for **(A)** DiR, **(B)** Cy5, and **(C)** GFP positive cell percentages for each LNP group. Source data are provided as a Supplementary Source Data file.

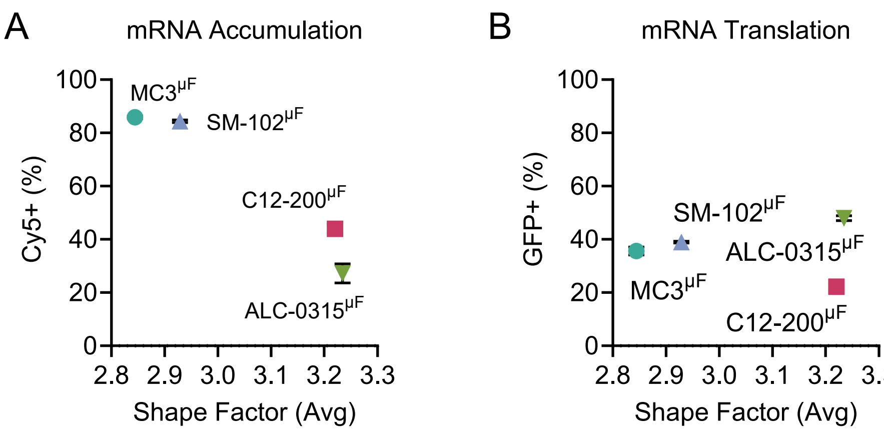


**Supplementary Figure 13: Shape factor vs T cell LNP uptake and transfection. A–B,** Plot of shape factor ($\frac{D_{max}}{R_g}$), determined by P(r) analyses of average peak SAXS profiles, against **(A)** Cy5 mRNA uptake and **(B)** GFP translation in human primary T cells obtained via flow cytometry. Only the microfluidic-formulated LNPs were compared.

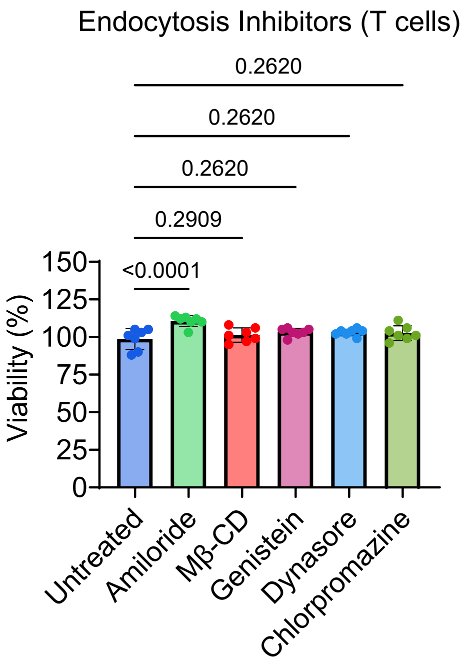


**Supplementary Figure 14: Viability of T cells with endocytosis inhibitors.** Human primary T cells (1:1, CD4+/CD8+) were activated for 16 h with human CD3/CD28 Dynabeads™ (1:1, bead/cell). Afterwards, the T cells were incubated with 2 mM amiloride, 500 μM methyl-β-cyclodextrin, 2 μM chlorpromazine, 100 μM Dynasore, and 200 μM genistein for 30 min. After another 24 h, viability was analyzed. Measurements are reported as mean ± SD of $n$ = 7 technical replicates. One-way ANOVA with *post hoc* Holm–Šídák correction for multiple comparisons was used to compare treated T cells with untreated T cells. Source data are provided as a Supplementary Source Data file.
